# Tight nanoscale clustering of Fcγ-receptors using DNA origami promotes phagocytosis

**DOI:** 10.1101/2021.03.18.436011

**Authors:** Nadja Kern, Rui Dong, Shawn M. Douglas, Ronald D. Vale, Meghan A. Morrissey

## Abstract

Macrophages destroy pathogens and diseased cells through Fcγ receptor (FcγR)-driven phagocytosis of antibody-opsonized targets. Phagocytosis requires activation of multiple FcγRs, but the mechanism controlling the threshold for response is unclear. We developed a DNA origami-based engulfment system that allows precise nanoscale control of the number and spacing of ligands. When the number of ligands remains constant, reducing ligand spacing from 17.5 nm to 7 nm potently enhances engulfment, primarily by increasing efficiency of the engulfment-initiation process. Tighter ligand clustering increases receptor phosphorylation, as well as proximal downstream signals. Increasing the number of signaling domains recruited to a single ligand-receptor complex was not sufficient to recapitulate this effect, indicating that clustering of multiple receptors is required. Our results suggest that macrophages use information about local ligand densities to make critical engulfment decisions, which has implications for the mechanism of antibody-mediated phagocytosis and the design of immunotherapies.

## Introduction

Immune cells eliminate pathogens and diseased cells while limiting damage to healthy cells. Macrophages, professional phagocytes and key effectors of the innate immune system, play an important role in this process by engulfing opsonized targets bearing ‘eat me’ signals. One of the most common ‘eat me’ signals is the immunoglobulin G (IgG) antibody, which can bind foreign proteins on infected cells or pathogens. IgG is recognized by Fcγ receptors (FcγR) in macrophages that drive antibody-dependent cellular phagocytosis (ADCP) (Dilillo, Tan, Palese, & Ravetch, 2014; Erwig & Gow, 2016; Nimmerjahn & Ravetch, 2008). ADCP is a key mechanism of action for several cancer immunotherapies including rituximab, trastuzumab, and cetuximab (Chao et al., 2010; Uchida et al., 2004; Watanabe et al., 1999; Weiskopf et al., 2013; Weiskopf & Weissman, 2015). Exploring the design parameters of effective antibodies could provide valuable insight into the molecular mechanisms driving ADCP.

Activation of multiple FcγRs is required for a macrophage to engulf a three-dimensional target. FcγR-IgG must be present across the entire target to drive progressive closure of the phagocytic cup that surrounds the target (Griffin, Griffin, Leider, & Silverstein, 1975). In addition, a critical antibody threshold across an entire target dictates an all-or-none engulfment response by the macrophage (Y. Zhang, Hoppe, & Swanson, 2010). Although the mechanism of this thresholded response remains unclear, receptor clustering plays a role in regulating digital responses in other immune cells (Berger et al., 2020; Davis & van der Merwe, 2006; Holowka & Baird, 1996; Kato et al., 2020; Ma, Lim, Benda, Goyette, & Gaus, 2020; Veneziano et al., 2020). FcγR clustering may also regulate phagocytosis (Goodridge, Underhill, & Touret, 2012). High resolution imaging of macrophages has demonstrated that IgG-bound FcγRs form clusters (resolution of >100 nm) within the plasma membrane (Lin et al., 2016; Lopes et al., 2017; Sobota et al., 2005). These small clusters, which recruit downstream effector proteins such as Syk-kinase and phosphoinositide 3-kinase, eventually coalesce into larger micron-scale patches as they migrate towards the center of the cell-target synapse (Jaumouillé et al., 2014; Lin et al., 2016; Lopes et al., 2017; Sobota et al., 2005).

Prior observational studies could not decouple ligand clustering from other parameters, such as ligand number or receptor mobility. As a result, we do not have a clear picture of how ligand number or molecular spacing regulate signal activation. To directly assess such questions, we have developed a reconstituted system that utilizes DNA origami to manipulate ligand patterns on a single-molecule level with nanometer resolution. We found that tightly spaced ligands strongly enhanced phagocytosis compared to the same number of more dispersed ligands. Through manipulating the number and spacing of ligands on individual origami pegboards, we found that 8 or more ligands per cluster maximized FcγR-driven engulfment, and that macrophages preferentially engulfed targets that had receptor-ligand clusters spaced ≤7 nm apart. We demonstrated that tight ligand clustering enhanced receptor phosphorylation, and the generation of PIP_3_ and actin filaments–critical downstream signaling molecules–at the phagocytic synapse. Together, our results suggest that the nanoscale clustering of receptors may allow macrophages to discriminate between lower density background stimuli and the higher density of ligands on opsonized targets. These results have implications for the design of immunotherapies that involve manipulating FcγR-driven engulfment.

## Results

### Developing a DNA-based chimeric antigen receptor to study phagocytosis

To study how isolated biochemical and biophysical ligand parameters affect engulfment, we sought to develop a well-defined and tunable engulfment system. Our lab previously developed a synthetic T cell signaling system, in which we replaced the receptor-ligand interaction (TCR-pMHC) with complimentary DNA oligos (Taylor, Husain, Gartner, Mayor, & Vale, 2017). We applied a similar DNA-based synthetic chimeric antigen receptor to study engulfment signaling in macrophages. In our DNA-CAR*γ* receptor, we replaced the native extracellular ligand binding domain of the Fcγ receptor with an extracellular SNAP-tag that covalently binds a benzyl-guanine-labeled single-stranded DNA (ssDNA) [receptor DNA; Figure 1a; (Morrissey et al., 2018)]. The SNAP-tag was then joined to the CD86 transmembrane domain followed by the intracellular signaling domain of the FcR γ chain (Nimmerjahn & Ravetch, 2008). We expressed the DNA-CARy in the macrophage-like cell line RAW264.7 and the monocyte-like cell line THP-1.

**Figure 1:**
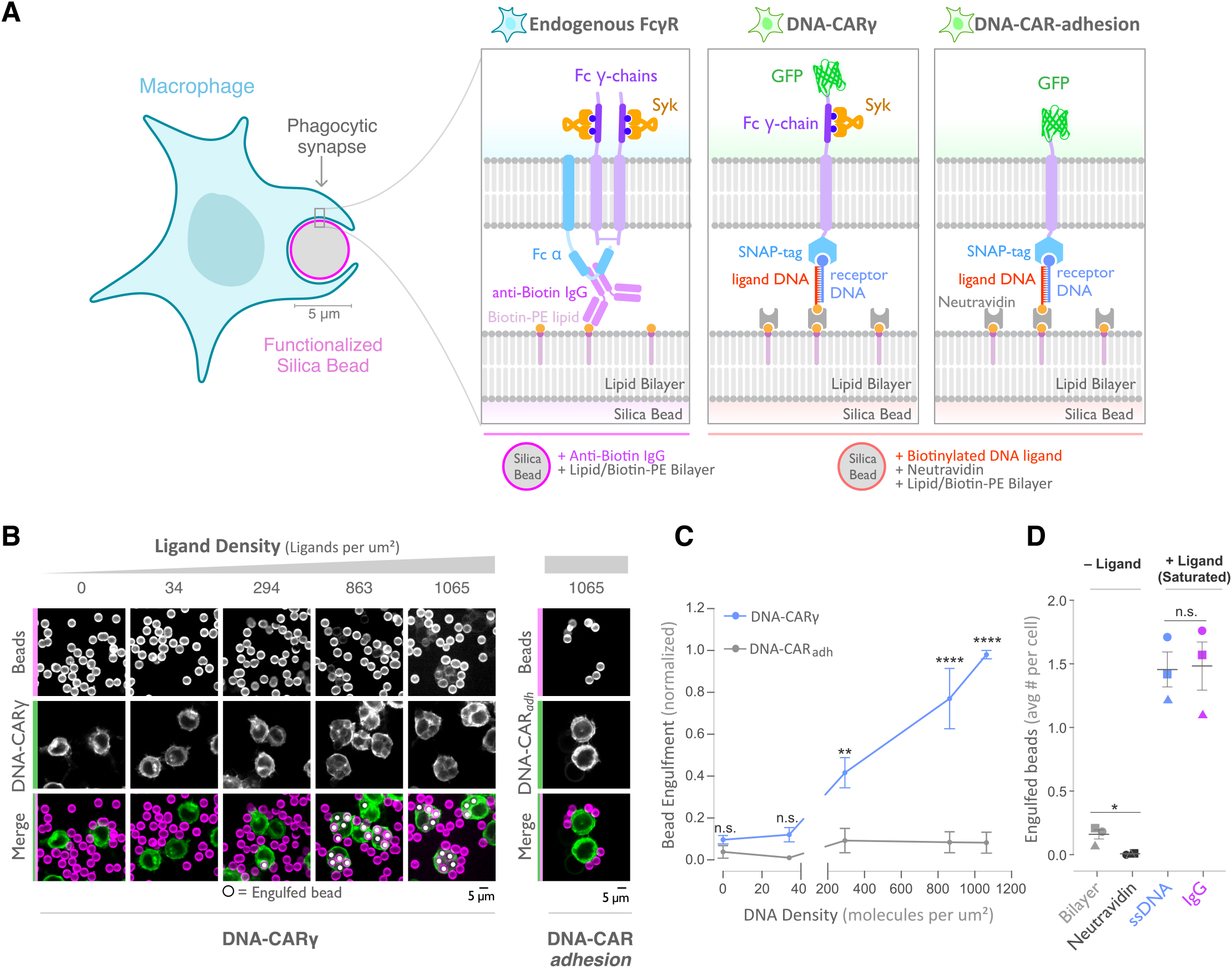
A DNA-based system for controlling engulfment. (A) Schematic shows the endogenous (left box) and DNA-based (middle and right boxes) engulfment systems. Engulfment via endogenous FcγRs (left box) is induced through anti-biotin IgG bound to 1-oleoyl-2-(12-biotinyl(aminododecanoyl))-sn-glycero-3-phosphoethanolamine (biotin-PE) lipids incorporated into the bilayer surrounding the silica bead targets. Engulfment induced via the DNA-based system uses chimeric antigen receptors (CAR) expressed in the macrophage and biotinylated ligand DNA that is bound to the lipid bilayer surrounding the silica bead. The DNA-CARy (middle box) consists of a ssDNA (receptor DNA) covalently attached to an extracellular SNAP-tag fused to a CD86 transmembrane domain, the intracellular domain of the FcR γ chain, and a fluorescent tag. The DNA-CAR_adhesion_ (right box) is identical but lacks the signaling FcR γ chain. (B) Example images depicting the engulfment assay. Silica beads were coated with a supported lipid bilayer (magenta) and functionalized with neutravidin and the indicated density of ligand DNA (Figure S1a). The functionalized beads were added to RAW264.7 macrophages expressing either the DNA-CAR*γ* or the DNA-CAR_adhesion_ (green) and fixed after 45 min. The average number of beads engulfed per macrophage was assessed by confocal microscopy. Scale bar denotes 5 µm here and in all subsequent figures. Internalized beads are denoted with a white sphere in the merged images. (C) The number of beads engulfed per cell for DNA-CAR*γ* (blue) or DNA-CAR_adhesion_ (grey) macrophages was normalized to the maximum bead eating observed in each replicate. Dots and error bars denote the mean ± SEM of three independent replicates (n≥100 cells analyzed per experiment). (D) DNA-CAR*γ* expressing macrophages were incubated with bilayer-coated beads (grey) functionalized with anti-biotin IgG (magenta), neutravidin (black), or neutravidin and saturating amounts of ssDNA (blue). The average number of beads engulfed per cell was assessed. Full data representing the fraction of macrophages engulfing specific numbers of IgG or ssDNA beads is shown in figure S1. Each data point represents the mean of an independent experiment, denoted by symbol shape, and bars denote the mean ± SEM. n.s. denotes p>0.05, * indicates p<0.05, ** indicates p<0.005 and **** indicates p<0.0001 by a multiple t-test comparison corrected for multiple comparisons using the Holm-Sidak method (C) or Student’s T-test (D).

As an engulfment target, we used silica beads coated with a supported lipid bilayer to mimic the surface of a target cell. The beads were functionalized with biotinylated ssDNA (ligand DNA) containing a sequence complementary to the receptor DNA via biotin-neutravidin interactions (Figure 1a). We used a ligand DNA strand that has 13 complementary base pairs to the receptor DNA, which we chose because the receptor-ligand dwell time (∼24 sec (Taylor et al., 2017)) was comparable to the dwell time of IgG-FcγR interactions (∼30-150 sec (Li et al., 2007)).

To test whether this synthetic system can drive specific engulfment of ligand-functionalized silica beads, we used confocal microscopy to measure the number of beads that were engulfed by each cell (Figure 1b, c). The DNA-CAR*γ* drove specific engulfment of DNA-bound beads in both RAW264.7 and THP-1 cells (Figure 1c, S1). The extent of engulfment was similar to IgG-coated beads, and the ligand density required for robust phagocytosis was also comparable to IgG [Figure 1d, S1; (Bakalar et al., 2018; Morrissey, Kern, & Vale, 2020)]. As a control, we tested a variant of the DNA-CAR that lacked the intracellular domain of the FcR γ chain (DNA-CAR_adhesion_). Cells expressing the DNA-CAR_adhesion_ failed to induce engulfment of DNA-functionalized beads (Figure 1c), demonstrating that this process depends upon the signaling domain of the FcγReceptor. Together, these data show that the DNA-CAR*γ* can drive engulfment of targets in a ligand- and FcγR-specific manner.

### DNA origami pegboards activate DNA-CAR*γ* macrophages

DNA origami technology provides the ability to easily build three-dimensional objects that present ssDNA oligonucleotides with defined nanometer-level spatial organization (Hong, Zhang, Liu, & Yan, 2017; Rothemund, 2006; Seeman, 2010; Shaw et al., 2019; Veneziano et al., 2020). We used DNA origami to manipulate the spatial distribution of DNA-CAR*γ* ligands in order to determine how nanoscale ligand spacing affects engulfment. We used a recently developed two-tiered DNA origami pegboard that encompasses a total of 72 ssDNA positions spaced 7 nm and 3.5 nm apart in the x and y dimensions, respectively (Dong et al., 2021)(Figure 2a, S2). Each of the 72 ligand positions can be manipulated independently, allowing for full control over the ligand at each position (Figure S2). The DNA origami pegboard also contains fluorophores at each of its four corners to allow for visualization, and 12 biotin-modified oligos on the bottom half of the pegboard to attach it to a neutravidin-containing supported lipid bilayer or glass coverslip (Figure 2a, b, S2).

**Figure 2:**
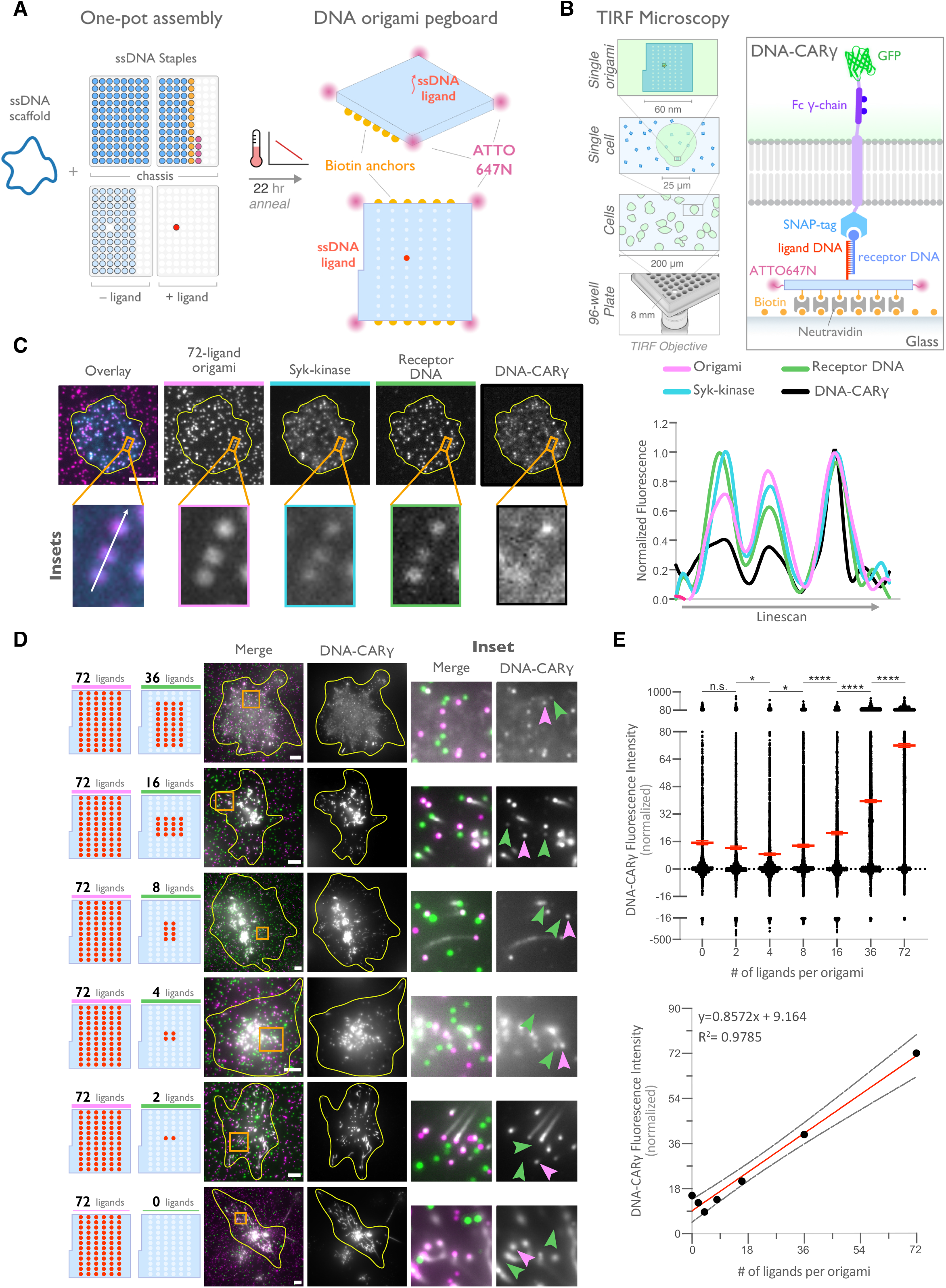
DNA origami pegboard induces ligand dependent signaling. (A) Schematic shows the DNA-origami pegboard used in this study (right) and the components used to create it using a one-pot assembly method (left, figure S2). The top of the two-tiered DNA origami pegboard has 72 positions spaced 7 nm and 3.5 nm apart in the x and y dimensions, which can be modified to expose a single-stranded ligand DNA (red) or no ligand (light blue). A fluorophore is attached at each corner of the pegboard for visualization (pink). The bottom tier of the pegboard displays 12 biotin molecules (yellow) used to attach the origami to neutravidin-coated surfaces. Full representation of the DNA origami pegboard assembly is shown in figure S2. (B) Schematic portraying the TIRF microscopy setup used to image THP-1 cells interacting with origami pegboards functionalized to glass coverslips in (C) and (D) (left). On the right is a zoomed-in side view of an origami pegboard functionalized to a biotin (yellow) and neutravidin (grey) functionalized glass coverslip and interacting with a single DNA-CAR*γ* receptor. (C) TIRF microscopy images of THP-1 cells show that the DNA-CAR*γ* (BFP; 5^th^ panel; black in linescan), the receptor DNA bound to the DNA-CAR*γ* (Cy5; 4^th^ panel; green in linescan), and Syk (mNeonGreen; 3^rd^ panel; cyan in merge and linescan) are recruited to individual 72-ligand origami pegboards (Atto-647; 2^nd^ panel; magenta in merge and linescan). Each diffraction limited magenta spot represents an origami pegboard. The top panels show a single cell (outlined in yellow), and the bottom insets (orange box in top image) show three origami pegboards at higher magnification. The linescan (right, area denoted with a white arrow in merged inset) shows the fluorescence intensity of each of these channels. Intensity was normalized so that 1 is the highest observed intensity and 0 is background for each channel. (D) TIRF microscopy images show DNA-CAR*γ* expressing THP1s interacting with 72-ligand origami pegboards (pink) and origami pegboards presenting the indicated number of ligands (pegboards labeled in green). Left schematics represent origami pegboard setups for each row of images where red dots denote the presence of a ligand DNA. Middle images depict a single macrophage (outlined in yellow), and right images show the area indicated with an orange box on the left. Examples of DNA-CAR*γ*-mNeonGreen (grey) recruitment to individual origami pegboards is marked by pink (72L origami pegboard) and green (origami pegboard with the indicated ligand number) arrowheads (right). (E) Quantification of experiment shown in (D). Top graph shows the DNA-CAR*γ* intensity at the indicated origami pegboard type normalized to the average DNA-CAR*γ* intensity at 72L origami pegboards in the same well. Each dot represents one origami pegboard and red lines denote the mean ± SEM of pooled data from three separate replicates. n.s. denotes p>0.05, * indicates p<0.05, and **** indicates p<0.0001 by an ordinary one-way ANOVA with Holm-Sidak’s multiple comparison test. A linear regression fit (bottom) of the average fluorescence intensities of each of the origami pegboards suggests that the mean DNA-CAR*γ* fluorescent intensities are linearly proportional to the number of ligands per DNA origami pegboard. The black dots represent the mean normalized DNA-CAR*γ* intensity, the red line denotes the linear regression fit, and the grey lines show the 95% confidence intervals.

To determine if the DNA origami pegboards could successfully activate signaling, we first tested whether receptors were recruited to the origami pegboard in a ligand-dependent manner. Using TIRF microscopy, we quantified the fluorescence intensity of the recruited GFP-tagged DNA-CAR*γ* receptor to origami pegboards presenting 0, 2, 4, 16, 36 or 72 ligands (Figure 2b-e). Using signal from the 72 ligand (72L) origami pegboard as an internal intensity standard of brightness, and thus correcting for differences in illumination between wells, we found that the average fluorescence intensity correlated with the number of ligands presented by individual origami pegboards (Figure 2d, e). In addition, we measured Syk recruitment to individual DNA origami pegboards and found that Syk intensity also increased as a function of the number of ligands present on each origami pegboard (Figure 2c, S3). These results confirmed that our DNA origami system provides a platform that allows quantitative receptor recruitment and the analysis of downstream signaling pathways.

### Nanoscale clustering of ligand enhances phagocytosis

FcγReceptors cluster upon ligand binding, but the functional importance of such clustering for phagocytosis has not been directly addressed, and whether a critical density of receptor-ligand pairs is necessary to initiate FcγR signaling is unclear (Duchemin, Ernst, & Anderson, 1994; Jaumouillé et al., 2014; Lin et al., 2016; Lopes et al., 2017; Sobota et al., 2005). To address these questions, we varied the size of ligand clusters by designing DNA origami pegboards presenting 2-36 ligands. To ensure a constant total number of ligands and origami pegboards on each bead, we mixed the signaling origami pegboards with 0-ligand “blank” origami pegboards in appropriate ratios (Figure 3a). We confirmed that the surface concentration of origami pegboards on the beads was comparable using fluorescence microscopy (Figure S4). We found that increasing the number of ligands per cluster increased engulfment, but that engulfment plateaued at a cluster size of 8 ligands (Figure 3b). We confirmed that the observed engulfment phenotype was both ligand, receptor, and FcγR signaling dependent (Figure 3c, d). Together, these data reveal that FcγReceptor clustering strongly enhances engulfment, up to a cluster size of 8 ligands.

**Figure 3:**
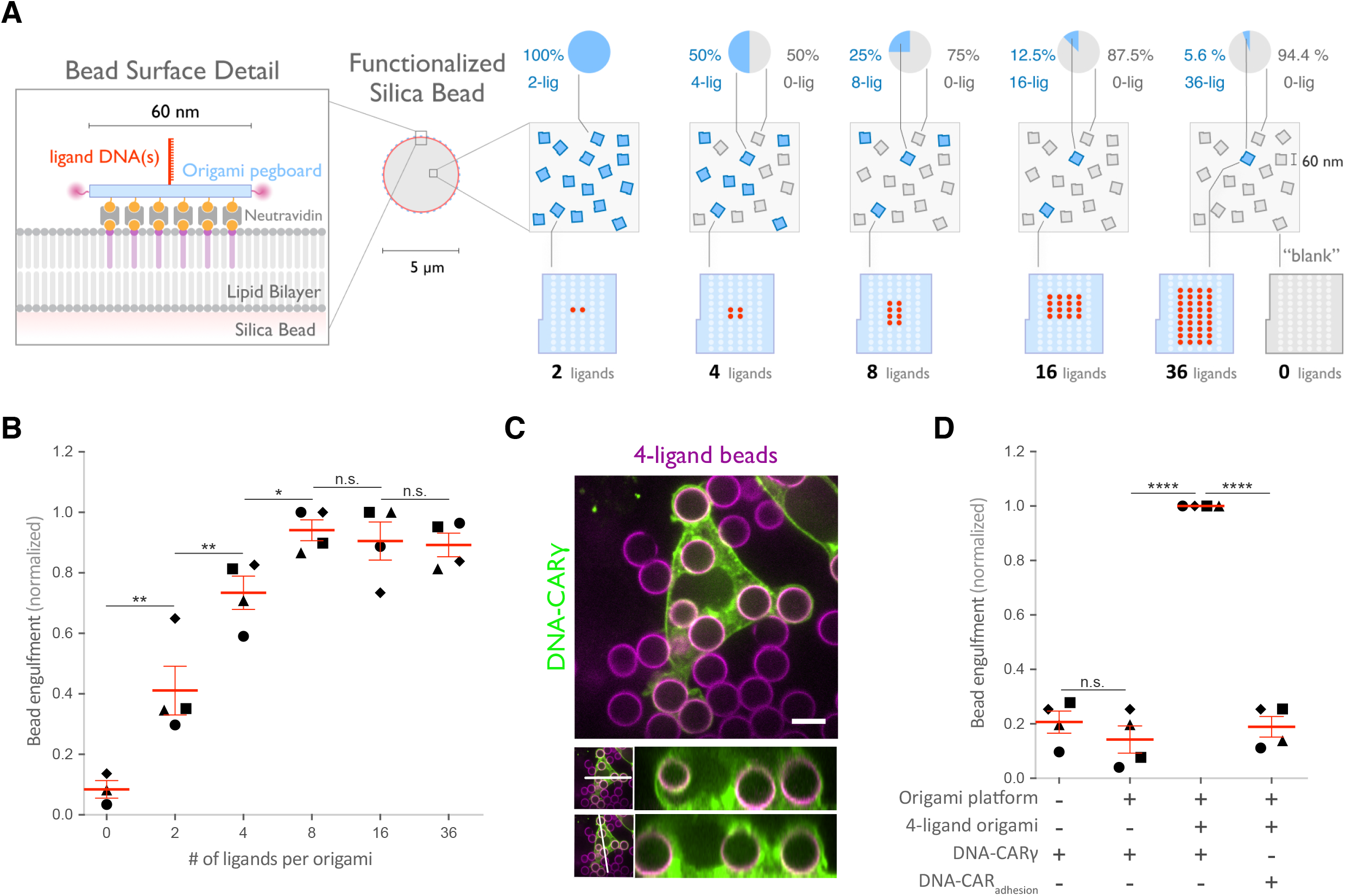
Nanoscale clustering of ligand enhances phagocytosis. (A) Schematic showing an origami pegboard functionalized to a lipid bilayer surrounding a silica bead (left) and the origami pegboard mixtures used to functionalize the bilayer-coated silica beads for experiment quantified in (B) (right). Blue squares represent origami pegboards with the indicated number of ligands (schematics below, red dot denotes ligand DNA and light blue dot denotes no ligand) and grey squares represent 0-ligand “blank” origami pegboards. Pie charts above describe the ratios of ligand origami presenting pegboards to “blank” pegboards. (B) Beads were functionalized with mixtures of origami pegboards containing the indicated ligand-presenting origami pegboard and the 0-ligand “blank” origami pegboards in amounts designated in (A). The graph depicts the number of beads internalized per DNA-CAR*γ* expressing macrophage normalized to the maximum bead eating in that replicate. Each dot represents an independent replicate (n≥100 cells analyzed per experiment), denoted by symbol shape, with red lines denoting mean ± SEM. Data is normalized to the maximum bead eating in each replicate. (C) Example image showing the DNA-CAR*γ* (green) drives engulfment of beads (bilayer labeled in magenta) functionalized with 4-ligand DNA origami pegboards. A cross section of the z plane indicated in the inset panel (white line, bottom), shows that beads are fully internalized. (D) Bilayer coated silica beads were functionalized with neutravidin, neutravidin and DNA origami pegboards presenting 0 DNA ligands, or neutravidin and 4-ligand DNA origami pegboards. The graph depicts normalized bead eating per cell of the indicated bead type for cells expressing the DNA-CAR*γ* or the DNA-CAR_adhesion_. Each dot represents an independent replicate, denoted by symbol shape (n≥100 cells analyzed per experiment), with red lines denoting mean ± SEM. The data are normalized to the maximum bead eating in each replicate. * denotes p<0.05, ** denotes p<0.005, **** denotes p<0.0001, and n.s. denotes p>0.05 in (B) and (D) as determined by an Ordinary one-way ANOVA with Holm-Sidak’s multiple comparison test.

### Spatial organization of ligands in nanoclusters regulates engulfment

Next, we examined whether distance between individual receptor-ligand molecules within a signaling cluster impacts engulfment. For this experiment, we varied the spacing of 4 ligands on the origami pegboard. The 4-ligand tight origami (4T) contains 4 ligands clustered at the center of the pegboard (7 nm by 3.5 nm square), the medium origami (4M) has ligands spaced 21 nm by 17.5 nm apart, and the spread origami (4S) has 4 ligands positioned at the four corners of the pegboard (35 nm by 38.5 nm square) (Figure 4a). We found that the efficiency of macrophage engulfment was approximately 2-fold higher for the 4T functionalized beads when compared to the 4M or 4S beads (Figure 4a). We confirmed via fluorescence microscopy that the concentration of origami pegboards on the surface was similar, and therefore ligand numbers on the beads were similar (Figure S5). DNA CAR constructs that have the FcR *γ* and ⍰ chain transmembrane domains in place of the CD86 transmembrane domain and human THP-1 cells expressing the DNA-CAR*γ* showed the same ligand spacing dependence (Figure S5). Expression of the various DNA CARs at the cell cortex was comparable, and engulfment of beads functionalized with both the 4T and the 4S origami platforms was dependent on the Fc*γ*R signaling domain (Figure S5). Together, these results demonstrate that macrophages preferentially engulf targets with tighter ligand clusters.

**Figure 4:**
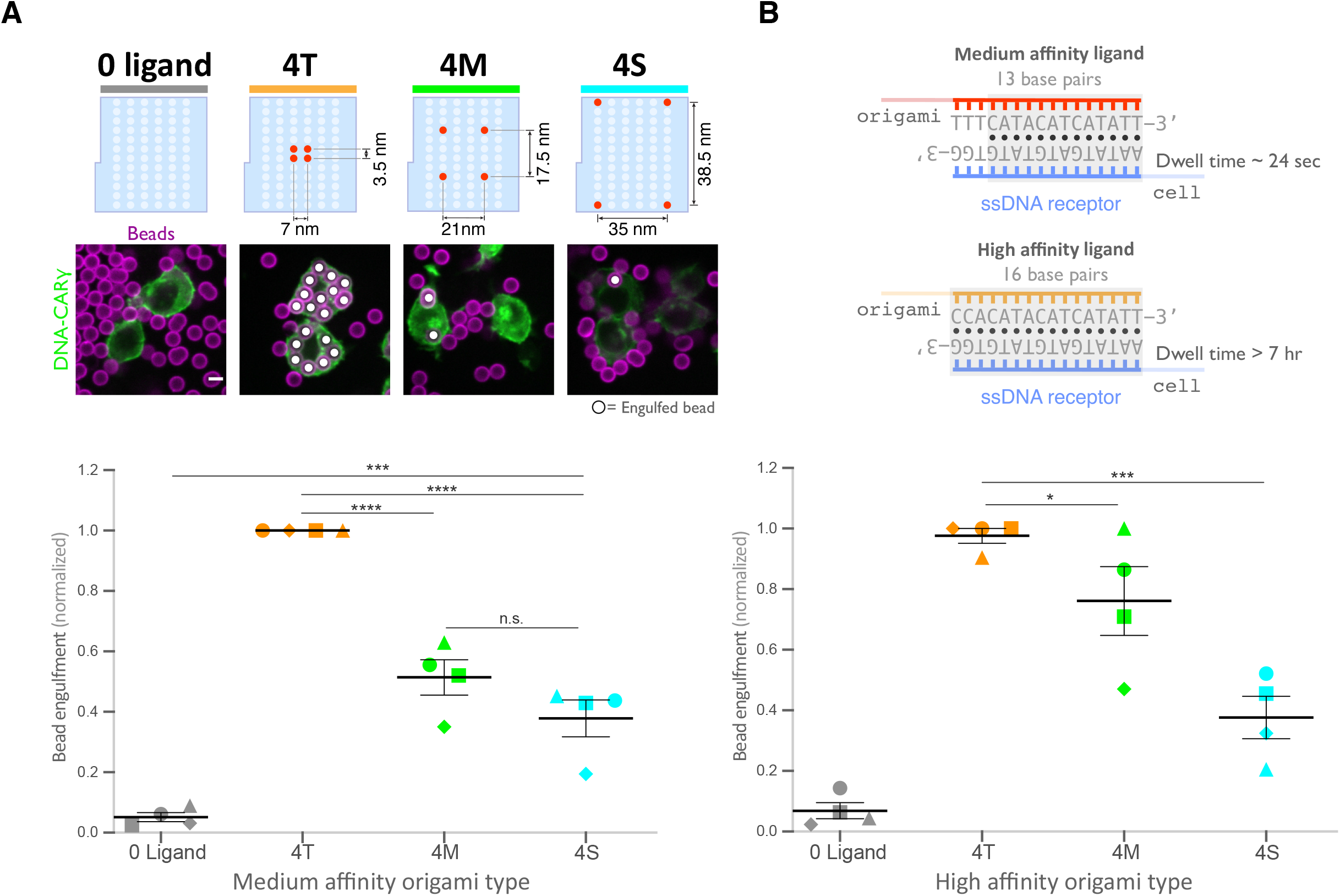
Spatial arrangement of ligands within nanoclusters regulates engulfment. (A) Schematics (top) depict 4-ligand origami pegboards presenting ligands at the positions indicated in red. Beads were functionalized with 0-ligand ‘blank’ (grey) origami pegboards, 4T (orange) origami pegboards, 4M (green) origami pegboards, or 4S (cyan) origami pegboards at equal amounts and fed to DNA-CAR*γ* expressing macrophages. Representative confocal images (middle) depict bead (bilayer in magenta) engulfment by macrophages (green). Internalized beads are denoted with a white sphere. Quantification of the engulfment assay is shown in the graph below depicting the number of beads engulfed per macrophage normalized to the maximum observed eating in that replicate. (B) Schematics of the receptor DNA (blue) paired with the medium affinity 13 base paired DNA-ligand (red) used in all previous experiments including (A) and the high affinity 16 base pair ligand-DNA (yellow) used for experiment shown in graph below. Beads were functionalized with 0-ligand ‘blank’ (grey), high affinity 4T (orange), high affinity 4M (green), or high affinity 4S (cyan) origami pegboards and fed to DNA-CAR*γ* expressing macrophages. Graph shows the number of beads engulfed per macrophage normalized to the maximum observed eating in that replicate. Each data point represents the mean of an independent experiment, shapes denote data from the same replicate, and bars show the mean ± SEM (A, B). * denotes p<0.05, *** denotes p<0.0005, **** denotes p<0.0001, and n.s. denotes p>0.05 as determined by an Ordinary one-way ANOVA with Holm-Sidak’s multiple comparison test (A, B).

Tightly spaced ligands could potentially increase phagocytosis by enhancing the avidity of receptor-ligand interactions within each cluster. Such a hypothesis would predict that tightly spaced ligands increase DNA-CAR*γ*-BFP occupancy at the phagocytic cup. However, when we measured the total fluorescence intensity of receptors at the phagocytic cup, we did not detect a difference in DNA-CAR*γ*-BFP recruitment to 4T and 4S beads (Figure 6a, b). However, to eliminate any potential contribution of avidity, we created 4T and 4S origami pegboards with very high-affinity 16mer DNA ligands that are predicted to dissociate on a time scale of >7 hr (Taylor et al., 2017) (Figure 4b). Using these 16mer high-affinity ligands, we found that 4T origami beads were still preferentially engulfed over 4M or 4S origami beads (Figure 4b, S5). These results suggest that an avidity effect is not the cause of the preferential engulfment of targets having tightly spaced ligands.

**Figure 6:**
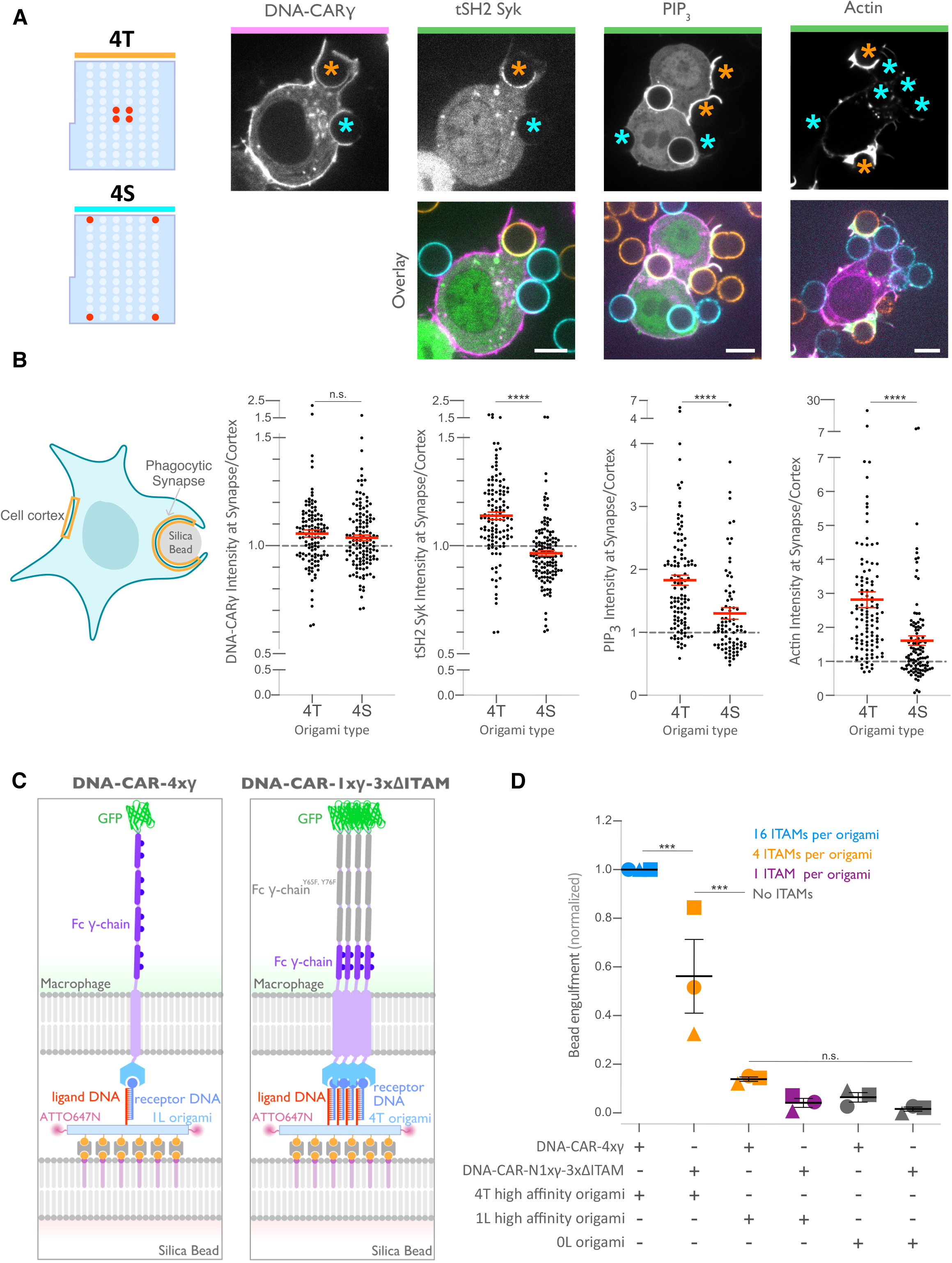
Nanoscale ligand spacing controls receptor activation. (A) Beads were functionalized with 4T (orange) or 4S (cyan) origami pegboards at equal amounts, added to macrophages expressing the DNA-CAR*γ* (magenta) and the indicated signaling reporter protein (green; greyscale on top). Phagocytic synapses were imaged via confocal microscopy. Asterisks indicate whether a 4T (orange) or a 4S (cyan) bead is at the indicated phagocytic synapse in the upper panel. (B) Schematic (left) depicts the areas measured from images shown in (A) to quantify the fluorescence intensity (yellow outlines). Each phagocytic synapse measurement was normalized to the fluorescence intensity of the cell cortex at the same z-plane. Graphs (right) depict the ratio of fluorescence at 4T or 4S functionalized bead synapses to the cortex for the indicated reporter. Each dot represents one bead with red lines denoting mean ± SEM. (C) Schematic portraying the CAR constructs and origami used in the experiment quantified in (D). The DNA-CAR-4xy construct (left) consists of four repeats of the intracellular domain of the DNA-CAR*γ* connected by a GGSG linker. The DNA-CAR-1xy-3xLlITAM (right) is identical to the DNA-CAR-4xy except that the tyrosines composing the ITAM domains (purple circles) are mutated to phenylalanines in the three C-terminal repeats (grey). Cells expressing either of these constructs were fed beads functionalized with either high affinity 1-ligand origami pegboards (left), high affinity 4T origami pegboards (right), or 0 ligand “blank” origami pegboards (not shown), and engulfment was assessed after 45 min. (D) Graph shows the number of beads engulfed per macrophage normalized to the maximum observed eating in that replicate. Each data point represents the mean from an independent experiment, denoted by symbol shape, and bars denote the mean ± SEM. Blue points represent a condition where 16 ITAMs are available per origami, orange points represent conditions where 4 ITAMs are available per origami, purple points represent a condition where 1 ITAM is available per origami, and grey points represent conditions where no ITAM is available. n.s. denotes p>0.05, *** denotes p<0.0005, and **** denotes p<0.00005 as determined by the Student’s T-test (B) or an Ordinary one-way ANOVA with Holm-Sidak’s multiple comparison test (D).

### Tight ligand spacing enhances engulfment initiation and downstream signaling

We next determined how ligand spacing affects the kinetics of engulfment. Using data from live-cell imaging, we subdivided the engulfment process into three steps: bead binding, engulfment initiation, and engulfment completion (Figure 5a, Supplemental movie 1). To compare engulfment dynamics mediated by 4T and 4S origami pegboards in the same experiment, we labeled each pegboard type with a different colored fluorophore, functionalized a set of beads with each type of pegboard, and added both bead types to macrophages at the same time (Figure 5b, Supplemental movie 2). Macrophages interacted with beads functionalized with the 4T and 4S pegboards with comparable frequency (46 ± 7% total bead-cell contacts vs. 54 ± 7% total bead-cell contacts respectively). However, the probability of engulfment initiation was significantly higher for the 4T (95 ± 5% of bead contacts) versus 4S (61 ± 9% of bead contacts) beads, and the probability that initiation events resulted in successful completion of engulfment was higher for 4T (69 ± 9% of initiation events) versus 4S (39 ± 11% of initiation events) beads (Figure 5a). Initiation events that failed to induce successful engulfment either stalled after progressing partially over the bead or retracted the extended membrane back to the base of the bead. In addition, for beads that were engulfed, the time from contact to engulfment initiation was ∼300 sec longer for beads functionalized with 4S origami pegboards than beads containing 4T origami pegboards (Figure 5c). However, once initiated, the time from initiation to completion of engulfment did not differ significantly for beads coated with 4T or 4S origami (Figure 5d). Overall, 66 ± 8% of 4T bead contacts resulted in successful engulfment compared to 24% ± 8% for 4S beads (Figure 5e). The DNA-CAR_adhesion_ macrophages rarely met the initiation criteria, suggesting that active signaling from the Fc*γ*R is required (Figure S6). Together, these data reveal that tighter spacing between ligands within a cluster enhances the probability and kinetics of initiating engulfment, as well as the overall success frequency of completing engulfment, but does not affect the rate of phagosome closure once initiated.

**Figure 5:**
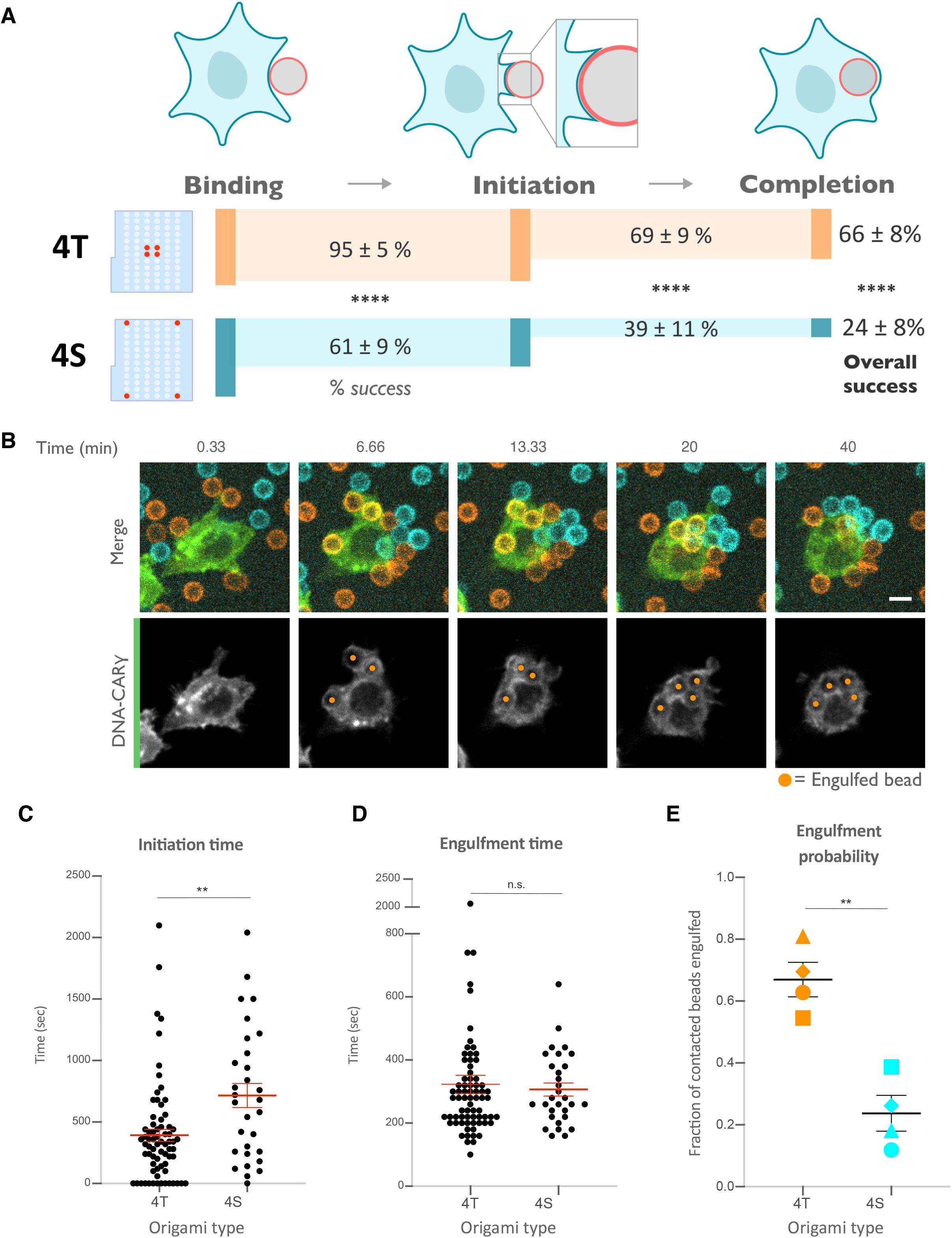
Nanoscale ligand clustering controls engulfment initiation. (A) Schematic portraying origami pegboards used to analyze the steps in the engulfment process quantified in (C), (D), and (E). Bead binding is defined as the first frame the macrophage contacts a bead; initiation is the first frame in which the macrophage membrane has begun to extend around the bead, and completion is defined as full internalization. The macrophage membrane was visualized using the DNA CARy, which was present throughout the cell cortex. The % of beads that progress to the next stage of engulfment (% success) is indicated for 4T (orange, origami labeled with Atto550N) and 4S (cyan, origami labeled with Atto647N) beads. **** denotes p<0.0001 as determined by Fischer’s exact test. (B) Still images from a confocal microscopy timelapse showing the macrophage (green) interacting with both the 4T origami pegboard functionalized beads (orange) and the 4S origami pegboard functionalized beads (cyan), but preferentially engulfing the 4T origami pegboard functionalized beads. In the bottom panel (DNA-CAR*γ* channel), engulfed beads have been indicated by a sphere colored to match its corresponding origami type. (C) Graph depicts quantification of the time from bead contact to engulfment initiation for all beads that were successfully engulfed. Each dot represents one bead with red lines denoting mean ± SEM. (D) Graph depicts the time from engulfment initiation to completion. Each dot represents one bead with red lines denoting mean ± SEM. (E) Graph shows the fraction of contacted 4T and 4S beads engulfed (orange and cyan, respectively) by the macrophages. Data represent quantification from 4 independent experiments, denoted by symbol shape, and bars denote the mean ± SEM. n.s. denotes p>0.05 and ** indicates p<0.005 by Student’s T-test comparing the 4T and 4S functionalized beads (C-E).

### Tightly spaced ligands enhance receptor phosphorylation

We next determined how the 4T or 4S origami pegboards affect signaling downstream of FcγR binding by measuring fold enrichment at the phagocytic cup compared to the rest of the cortex of 1) a marker for receptor phosphorylation (the tandem SH2 domains of Syk, (Bakalar et al., 2018; Morrissey et al., 2018)), 2) PIP_3_ (via recruitment of the PIP_3_ binding protein Akt-PH-GFP), and 3) filamentous actin (measured by rhodamine-Phalloidin binding, Figure 6a, b). We found that 4T phagocytic cups recruited more tSH2-Syk than the 4S beads, indicating an increase in receptor phosphorylation by nano-clustered ligands. Generation of PIP_3_ and actin filaments at the phagocytic cup also increased at 4T relative to 4S synapses (Figure 6b). This differential recruitment of downstream signaling molecules to 4T versus 4S origami beads was most apparent in early and mid-stage phagocytic cups; late-stage cups showed only a slightly significant difference in tSH2-Syk recruitment and no significant differences in generation of PIP_3_ or actin filaments (Figure S7). Together, these data demonstrate that nanoscale ligand spacing affects early downstream signaling events involved in phagocytic cup formation.

We next sought to understand why distributing ligands into tight clusters enhanced receptor phosphorylation and engulfment. One possibility is that the clustering of four complete receptors is needed to drive segregation of the inhibitory phosphatase CD45 and allow sustained phosphorylation of the Fc*γ*R Immune Receptor Tyrosine-based Activation Motif (ITAM) (Bakalar et al., 2018; Freeman et al., 2016; Goodridge et al., 2012; Schmid et al., 2016). Alternatively, the 4-ligand cluster may be needed to obtain a critical intracellular concentration of Fc*γ*R ITAM signaling domains. To test for the latter possibility, we designed a synthetic receptor (DNA-CAR-4x*γ*) that contains four repeats of the intracellular domain of the DNA-CAR*γ* connected by a GGSG linker between each repeat (Figure 6c). We confirmed that this DNA-CAR-4x*γ* receptor was more potent in activating engulfment than an equivalent receptor (DNA-CAR-1x*γ*-3x*Δ*lITAM) in which the 3 C-terminal ITAM domains were mutated to phenylalanines (Figure 6c, d). Keeping the number of intracellular ITAMs constant, we compared the engulfment efficiency mediated by two different receptors: 1) the DNA-CAR-4x*γ* that interacted with beads functionalized with 1-ligand origami, and 2) the DNA-CAR-1x*γ*-3x*Δ*lITAM that interacted with beads coated with equivalent amounts of 4T origami (Figure 6c). While the DNA-CAR-1x*γ*-3x*Δ*lITAM-expressing macrophages engulfed 4T origami beads, the DNA-CAR-4x*γ* macrophages failed to engulf the high-affinity 1-ligand origami beads (Figure 6d, Figure S7). To ensure that all four ITAM domains on the DNA-CAR-4x*γ* were signaling competent, we designed two additional DNA CARs which placed the functional ITAM at the second and fourth position (Figure S7). These receptors were able to induce phagocytosis of 4T origami beads, indicating that the DNA-CAR-4x*γ* likely contains 4 functional ITAMs. Collectively, these results indicate that the tight clustering of multiple receptors is necessary for engulfment and increasing the number of intracellular signaling modules on a single receptor is not sufficient to surpass the threshold for activation of engulfment.

## Discussion

Macrophages integrate information from many FcγR-antibody interactions to discriminate between highly opsonized targets and background signal from soluble antibody or sparsely opsonized targets. How the macrophage integrates signals from multiple FcγR binding events to make an all-or-none engulfment response is not clear. Here, we use DNA origami nanostructures to manipulate and assess how the nanoscale spatial organization of receptor-ligand interactions modulates FcγR signaling and the engulfment process. We found that tight ligand clustering increases the probability of initiating phagocytosis by enhancing FcγR phosphorylation.

Phagocytosis requires IgG across the entire target surface to initiate local receptor activation and to ‘zipper’ close the phagocytic cup (Freeman et al., 2016; Griffin et al., 1975). Consistent with this zipper model, incomplete opsonization of a target surface, or micron-scale spaces between IgG patches, decreases engulfment (Freeman et al., 2016; Griffin et al., 1975). Initially suggested as an alternative to the zipper model, the trigger model proposed that engulfment occurs once a threshold number of receptors interact with IgG (Ben M’Barek et al., 2015; Griffin et al., 1975; Swanson & Baer, 1995). While this model has largely fallen out of favor, more recent studies have found that a critical IgG threshold is needed to activate the final stages of phagocytosis (Y. Zhang et al., 2010). Our data suggest that there may also be a nanoscale density-dependent trigger for receptor phosphorylation and downstream signaling. Taken together, these results suggest that both tight nanoscale IgG-FcγR clustering and a uniform distribution of IgG across the target are needed to direct signaling to ‘zipper’ close the phagocytic cup. Why might macrophages use this local density dependent trigger to dictate engulfment responses? Macrophages constantly encounter background “eat me” signals (Gonzalez-Quintela et al., 2008). This hyper-local density measurement may buffer macrophages against background stimuli and weakly opsonized targets that are unlikely to have adjacent bound antibodies, while still robustly detecting and efficiently engulfing highly-opsonized targets.

Our findings are consistent with previous results demonstrating that FcγR crosslinking correlates with increased ITAM phosphorylation (M. M. Huang et al., 1992; Kwiatkowska & Sobota, 2001; Lin et al., 2016; Sobota et al., 2005). While our data pinpoints a role for ligand spacing in regulating receptor phosphorylation, it is possible that later steps in the phagocytic signaling pathway are also directly affected by ligand spacing. The mechanism by which dense-ligand clustering promotes receptor phosphorylation remains an open question, although our data rule out a couple of models. Specifically, we demonstrate that nanoscale ligand clustering does not noticeably affect the amount of ligand-bound receptor at the phagocytic cup, and that ligand spacing continues to affect engulfment when avidity effects are diminished through the use of high affinity receptor-ligands. Collectively, these data reveal that changes in receptor binding or recruitment caused by increased avidity are unlikely to account for the increased potency of clustered ligands. Our data also exclude the possibility that receptor clustering simply increases the local intracellular concentration of FcγR signaling domains, as arranging FcγR ITAMs in tandem did not have the same effect as clustering multiple receptor-ligand interactions. However, it remains possible that the geometry of the intracellular signaling domains could be important for activating or localizing downstream signaling, and that tandem ITAMs on the same polypeptide cannot produce the same engulfment signals as ITAMs on separate parallel polypeptides.

One possible model to explain the observed ligand-density dependence of signaling involves the ordering of lipids around the FcγReceptor. Segregated liquid-ordered and liquid-disordered membrane domains around immune receptor clusters have been reported to promote receptor phosphorylation (Bag, Wagenknecht-Wiesner, Lee, Shi, & Holowka, 2020; Dinic, Riehl, Adler, & Parmryd, 2015; Eggeling et al., 2009; Kabouridis, 2006; Simons & Ikonen, 1997; Sohn, Tolar, Jin, & Pierce, 2006; Stone, Shelby, Nnñez, Wisser, & Veatch, 2017). FcγR clusters are associated with liquid-ordered domains (Beekman, van der Linden, van de Winkel, & Leusen, 2008; Katsumata et al., 2001; Kwiatkowska & Sobota, 2001). Liquid-ordered domains recruit Src family kinases, which phosphorylate FcγRs, while liquid-disordered domains are enriched in the transmembrane phosphatase CD45, which dephosphorylates FcγRs (Bag et al., 2020; Sohn et al., 2006; Stone et al., 2017). Thus, lipid ordering could provide a mechanism that leads to receptor activation if denser receptor-ligand clusters are more efficient in nucleating or associating with ordered lipid domains.

As an alternative model, a denser cluster of ligated receptors may enhance the steric exclusion of the bulky transmembrane proteins like the phosphatases CD45 and CD148 (Bakalar et al., 2018; Goodridge et al., 2012; Zhu, Brdicka, Katsumoto, Lin, & Weiss, 2008). CD45 is heavily glycosylated, making the extracellular domain 25-40 nm tall (Davis & van der Merwe, 2006; McCall, Shotton, & Barclay, 1992; Woollett, Williams, & Shotton, 1985). Because of its size, CD45 is excluded from close cell-cell contacts, such as those mediated by IgG-FcγR, which have a dimension of 11.5 nm (Bakalar et al., 2018; Burroughs et al., 2011; Carbone et al., 2017; Chung, Koo, & Boxer, 2013; Lu, Ellsworth, Hamacher, Oak, & Sun, 2011; Schmid et al., 2016). IgG bound to antigens ≤10.5 nm from the target surface induces CD45 exclusion and engulfment (estimated total intermembrane distance of ≤22 nm (Bakalar et al., 2018)). Our DNA origami structure is estimated to generate similar intermembrane spacing, consisting of hybridized receptor-ligand DNA (∼9.4 nm), the origami pegboard (6 nm) and neutravidin (4 nm) (Rosano, Arosio, & Bolognesi, 1999)]. A higher receptor-ligand density constrains membrane shape fluctuations (Krobath, Rózycki, Lipowsky, & Weikl, 2009, 2011; Rózycki, Lipowsky, & Weikl, 2010), and this constraint may increase CD45 exclusion (Schmid et al., 2016). Both the lipid ordering and the steric exclusion models predict at least a partial exclusion of the CD45 from the zone of the receptor cluster. However, the dimension of the tight cluster in particular is very small (7 by 3.5 nm) and measurement of protein concentration at this level is currently not easily achieved, even with super-resolution techniques. Overall, our results establish the molecular and spatial parameters necessary for FcγR activation and demonstrate that the spatial organization of IgG-FcγR interactions alone can affect engulfment decisions.

How does the spacing requirements for Fc*γ*R nanoclusters compare to other signaling systems? Engineered multivalent Fc oligomers revealed that IgE ligand geometry alters Fcε receptor signaling in mast cells (Sil, Lee, Luo, Holowka, & Baird, 2007). DNA origami nanoparticles and planar nanolithography arrays have previously examined optimal inter-ligand distance for the T cell receptor, B cell receptor, NK cell receptor CD16, death receptor Fas, and integrins (Arnold et al., 2004; Berger et al., 2020; Cai et al., 2018; Deeg et al., 2013; Delcassian et al., 2013; Dong et al., 2021; Veneziano et al., 2020). Some systems, like integrin-mediated cell adhesion, appear to have very discrete threshold requirements for ligand spacing while others, like T cell activation, appear to continuously improve with reduced intermolecular spacing(Arnold et al., 2004; Cai et al., 2018). Our system may be more similar to the continuous improvement observed in T cell activation, as our most spaced ligands (36.5 nm) are capable of activating some phagocytosis, albeit not as potently as the 4T. Interestingly, as the intermembrane distance between T cell and target increases, the requirement for tight ligand spacing becomes more stringent (Cai et al., 2018). This suggests that IgG bound to tall antigens may be more dependent on tight nanocluster spacing than short antigens. Planar arrays have also been used to vary inter-cluster spacing, in addition to inter-ligand spacing (Cai et al., 2018; Freeman et al., 2016). Examining the optimal inter-cluster spacing during phagosome closure may be an interesting direction for future studies.

Our study on the spatial requirements of FcγR activation could have implications for the design of therapeutic antibodies or chimeric antigen receptors. Antibody therapies that rely on FcγR engagement are used to treat cancer, autoimmune and neurodegenerative diseases (Chao et al., 2010; Nimmerjahn & Ravetch, 2005; Uchida et al., 2004; Watanabe et al., 1999; Weiskopf et al., 2013; Weiskopf & Weissman, 2015). Multimerizing Fc domains, or targeting multiple antibodies to the same antigen may increase antibody potency (X. Zhang et al., 2016). Interestingly, Rituximab, a successful anti-CD20 therapy that potently induces ADCP, has two binding sites on its target antigen (Zhao et al., 2020). Selecting clustered antigens, or pharmacologically inducing antigen clustering may also increase antibody potency (Chew et al., 2020). These results suggest that oligomerization may lead to more effective therapy; however, a systematic study of the spatial parameters that affect FcγR activation has not been undertaken (Bakalar et al., 2018). Our data suggest that antibody engineering strategies that optimize spacing of multiple antibodies through leucine zippers, cysteine bonds, DNA hybridization (Delcassian et al., 2013; Seifert et al., 2014; Sil, Lee, Luo, Holowka, & Baird, 2007) or multimeric scaffolds (Divine et al., 2020; Fallas et al., 2017; X. Huang et al., 2020; Ueda et al., 2020) could lead to stronger FcγR activation and potentially more effective therapies.

## Supporting information

Key Resources

## Supplemental Figure Legends

**Figure S1, related to Figure 1: DNA-based engulfment system reflects endogenous engulfment**

(A) Graph depicts the calibration used to determine the surface density of ssDNA on beads used in Figure 1b, c. The intensity of Alexa Fluor 647 fluorescent bead standards (black dots) was measured, and a simple linear regression (red line) was fit to the data. The fluorescence intensity of Alexa Fluor 647-ssDNA coated beads (blue dots) was measured, and the surface density was interpolated using the regression determined from the fluorescent bead standards. The concentration of ssDNA used for each bead coupling condition is indicated next to the blue points on the graph. (B) Macrophages expressing the DNA-CAR*γ* (blue) or the DNA-CAR_adhesion_ (grey) engulfed similar distributions of IgG functionalized beads. Data is pooled from two independent replicates. (C) Graph depicts the fraction of macrophages engulfing the indicated number of IgG (magenta) or ssDNA (blue) beads from data pooled from the three independent replicates presented in Figure 1d. (D) Graph shows the average number of Neutravidin (black), ligand-DNA (blue), or IgG (magenta) functionalized beads engulfed by the monocyte-like cell line THP1. Lines denote the mean engulfment from each independent replicate and bars denote ± SEM. P values were calculated using the Mann-Whitney test (B, C) and n.s. denotes p>0.05 as determined by the Student’s T-test (D).

**Figure S2, related to Figure 2: Design and Assembly of Nanoscale Ligand-Patterning Pegboard built from DNA origami**.

(A) 2D schematic of origami scaffold and staples. The p8064 ssDNA scaffold is combined with 160 ssDNA staples that form the chassis, biotin-modified surface anchors, and ATTO647N-labeled dyes, plus a combination of 72 ligand-patterning staples. We used three variants of the ligand-patterning staples: “-ligand” that lacks a 3’ single-stranded overhang and terminates flush with the pegboard surface, and a “medium-affinity” (red) and “high-affinity” (yellow) that form 13-bp and 16-bp duplexes with the DNA-CAR receptors, respectively. Assembly is performed by thermal annealing in a one-pot reaction. (B) Cadnano strand diagram for the pegboard with 72 medium-affinity ligands included. (C) Fourteen pegboard configurations were used in this study. Configurations are labeled by ligand count, spacing, and ligand affinity, and the corresponding plate wells used in each assembly are shown.

**Figure S3, related to Figure 2: Syk intensity increases with ligand number in origami cluster**

(A) TIRF microscopy images showing DNA-CAR*γ*-mNeonGreen and Syk-BFP expressing THP1s interacting with 72-ligand origami pegboards (pink) and origami pegboards presenting the indicated number of ligands (green) plated together on a glass surface (schematics shown on the left). Middle images depict a single macrophage, and right images show the area indicated with a yellow box on the left. Examples of Syk-BFP (grey) recruitment to individual origami pegboards is marked by pink (72L origami) and green (indicated ligand number origami) arrowheads (right). (B) Top graph shows the Syk intensity at each indicated origami pegboard type normalized to the average Syk intensity at 72L origami pegboards for each condition. Each dot represents the normalized Syk intensity at one origami and red lines denote the mean ± SEM of pooled data from three separate replicates. At ligand numbers fewer than 16, we did not detect Syk enrichment over background fluorescence of cytosolic Syk. A linear regression fit (bottom) of the average Syk fluorescence intensity at each origami pegboard type suggests that the mean Syk recruitment is linearly proportional to the number of ligands per DNA origami. n.s. denotes p>0.05 and **** indicates p<0.0001 by an ordinary one-way ANOVA with Holm-Sidak’s multiple comparison test.

**Figure S4, related to Figure 3: Origami intensity on beads is comparable across conditions**

(A) Graph shows the average Atto647N fluorescence intensity from the beads used in Figure 3a, b measured using confocal microscopy. Each dot represents an independent replicate (n≥100 cells analyzed per experiment), denoted by symbol shape, with red lines denoting mean ± SEM. n.s. denotes p>0.05 as determined by an Ordinary one-way ANOVA with Holm-Sidak’s multiple comparison test.

**Figure S5, related to Figure 4: Ligand clustering enhances engulfment in RAW macrophages expressing DNA CARs with endogenous Fc**Y**R transmembrane domains and in THP1s**

(A) Graph shows the average Atto647N fluorescence intensity from the beads used in Figure 4a measured using confocal microscopy. (B) Beads were functionalized with the indicated ligand-presenting origami pegboards in amounts calculated to equalize the total number of origami pegboards and ligands across conditions. Schematics (left) depict the origami utilized, where the positions presenting a ligand (red dots) and the positions not occupied by a ligand (light blue) are indicated. Graph (right) depicts the average number of the indicated type of beads internalized per DNA-CAR*γ*-expressing THP1, normalized to the maximum bead eating in that replicate. (C) Graph shows the average Atto647N647 fluorescence intensity from the beads used in Figure 4b measured using confocal microscopy. (D) Schematics below graph depict the DNA CAR constructs designed with varying transmembrane domains. Beads were functionalized with 4T origami pegboards (orange), 4S origami pegboards (cyan), or 0-ligand ‘blank’ origami pegboards (grey) and fed to macrophages expressing the DNA CAR receptor depicted below each section of the graph. Graph depicts the number of beads engulfed per macrophage normalized to the maximum observed eating in that replicate. (E) Graph shows the average Atto647N fluorescence intensity from the beads used in (D) measured using confocal microscopy. (F) DNA CAR receptors used in (D) are expressed and trafficked to the membrane at similar levels. Fluorescent intensity at the cell cortex of the DNA CAR-infected macrophage was quantified using the mean intensity of a 2 pixel width linescan at the cell membrane, with the mean intensity of a linescan immediately adjacent to the cell subtracted for local background. The fluorescence intensity was normalized to the average intensity of the DNA CAR_adhesion_ in each experiment. Each dot represents an individual cell and data is pooled from 3 independent experiments, with red lines denoting mean ± SEM. n.s. denotes p>0.05, * denotes p<0.05, ** denotes p<0.005, *** denotes p<0.0005, and **** indicates p<0.0001 as determined by an Ordinary one-way ANOVA with Holm-Sidak’s multiple comparison test (A-F).

**Figure S6, related to Figure 5: DNA CAR**_**adhesion**_ **fails to induce frequent engulfment initiation attempts**

(A) The average number of 4T origami pegboard-functionalized beads contacting (grey), in the initiation stage of engulfment (blue), or fully engulfed (green) by macrophages expressing either the DNA CAR_adhesion_ or the DNA CAR*γ* were quantified from fixed still images after 45 minutes of engulfment. 125 beads in contact with DNA CAR expressing macrophages were analyzed in 3 independent replicates. Bars represent the average number of beads identified at each stage and black lines denote ± SEM between replicates. n.s. denotes p>0.05 and * denotes p<0.05 as determined by an unpaired t-test with Holm-Sidak’s multiple comparison test.

**Figure S7, related to Figure 6: Differential recruitment of downstream signaling molecules is greater at early and mid-stage phagocytic cups**

(A) Data from experiment shown in Figure 6b is separated by early (macrophage membrane extends across <30% of the bead, left), mid (macrophage membrane extends across 30-70% of the bead, middle), and late (macrophage membrane extends across >70% of the bead, right) stage phagocytic cups. Graphs depict the ratio of fluorescence intensity at 4T or 4S functionalized bead synapses compared to the cortex. Each dot represents one bead with red lines denoting mean ± SEM. n.s. denotes p>0.05, * denotes p<0.05, *** denotes p<0.0005, and **** denotes p<0.00005 by the Student’s T-test. (B) Graph shows the average Atto647N fluorescence intensity from the beads used in Figure 6d measured using confocal microscopy. (C) Schematics depict the DNA-CAR-4x*γ* constructs used for experiment quantified in (D). (D) DNA CAR constructs shown in (C) were expressed in RAW macrophages and fed beads functionalized with 4T high affinity origami pegboards, 1 ligand high affinity origami pegboards, or 0 ligand origami pegboards. Graph depicts the number of beads engulfed per macrophage normalized to the maximum observed eating in that replicate. Each data point represents the mean from an independent experiment, denoted by symbol shape, and bars denote the mean ± SEM. Blue points represent a condition where 16 ITAMs are available per origami, orange points represent conditions where 4 ITAMs are available per origami, purple points represent a condition where 1 ITAM is available per origami, and grey points represent conditions where no ITAM is available. (E) Graph shows the average Atto647N fluorescence intensity from the beads used in (D) measured using confocal microscopy. (F) DNA CAR receptors used in (D) are expressed and trafficked to the membrane at similar levels. Fluorescent intensity at the cell cortex of the DNA CAR infected macrophage was quantified using the mean intensity of a 2 pixel width linescan at the cell membrane, with the mean intensity of a linescan immediately adjacent to the cell subtracted for local background. The fluorescence intensity was normalized to the average intensity of the DNA-CAR-4x*γ* in each experiment. Each dot represents an individual cell and data is pooled from 3 independent experiments, with red lines denoting mean ± SEM. n.s. denotes p>0.05 and **** indicates p<0.0001 as determined by an Ordinary one-way ANOVA with Holm-Sidak’s multiple comparison test (B,D-F).

**Supplemental movie 1: The engulfment program broken into three steps, bead binding, engulfment initiation, and engulfment completion**.

A macrophage infected with the DNA-CAR*γ* (green) engulfs a 5 um silica bead coated in a supported lipid bilayer (magenta) and functionalized with 4T origami pegboards. The movie is a maximum intensity projection of z-planes and depicts the bead binding, initiation, and completion steps of the engulfment process. Time is indicated at the top left and scale bar denotes 5 um.

**Supplemental movie 2: DNA CAR*γ* macrophages preferentially engulf beads functionalized with tightly spaced ligands**.

A DNA-CAR*γ* expressing macrophage (green) interacts with 4T origami pegboard functionalized beads (orange) and 4S origami pegboard functionalized beads (cyan) that were added simultaneously and in equal amounts to the well of cells. The macrophage engulfs only 4T origami pegboard functionalized beads. The movie is a maximum intensity projection of z-planes acquired every 20 secs for 28 min. Time is indicated at the top left.

## Methods

### Cell culture

RAW264.7 macrophages were purchased from the ATCC and cultured in DMEM (Gibco, Catalog #11965–092) supplemented with 1x Penicillin-Streptomycin-L-Glutamine (Corning, Catalog #30–009 Cl), 1 mM sodium pyruvate (Gibco, Catalog #11360-070) and 10% heat-inactivated fetal bovine serum (Atlanta Biologicals, Catalog #S11150H). THP1 cells were also purchased from the ATCC and cultured in RPMI 1640 Medium (Gibco, Catalog #11875-093) supplemented with 1x Pen-Strep-Glutamine and 10% heat-inactivated fetal bovine serum. All cells were certified mycoplasma-free and discarded after 20 passages to minimize variation.

### Constructs and antibodies

All relevant information can be found in the key resources table, including detailed descriptions of the amino acid sequences for all constructs.

### Lentivirus production and infection

Lentiviral infection was used to express constructs described in the key resources table in either RAW264.7 or THP1 cells. Lentivirus was produced by HEK293T cells or Lenti-X 293T cells (Takara Biosciences, Catalog #632180) transfected with pMD2.G (a gift from Didier Tronon, Addgene plasmid # 12259 containing the VSV-G envelope protein), pCMV-dR8.91 (since replaced by second generation compatible pCMV-dR8.2, Addgene plasmid #8455), and a lentiviral backbone vector containing the construct of interest (derived from pHRSIN-CSGW, see key resource table) using lipofectamine LTX (Invitrogen, Catalog # 15338–100). The HEK293T media was harvested 60-72 hr post-transfection, filtered through a 0.45 µm filter, and concentrated using Lenti-X (Takara Biosciences, Catalog #631232) via the standard protocol. Concentrated virus was added directly to the cells and the plate was centrifuged at 2200xg for 45 min at 37°C. Cells were analyzed a minimum of 60 hr later. Cells infected with more than one viral construct were FACs sorted (Sony SH800) before use to enrich for double infected cells.

### DNA origami preparation

The DNA origami pegboard utilized for all experiments was generated as described in figure S2. The p8064 DNA scaffold was purchased from IDT (Catalog # 1081314). All unmodified oligonucleotides utilized for the origami were purchased from IDT in 96 well plates with standard desalting purification and resuspension at 100 µM in water. Fluorophore and biotin conjugated oligonucleotides were also purchased from IDT (HPLC purification). All oligonucleotide sequences are listed in table 1, the assembly is schematized in figure S2, and the Cadnano strand diagram for the pegboard with 72 medium-affinity ligands is included in S2. Core staple oligonucleotides (200 nM) (plates 1 and 2), ligand oligonucleotides (200nM) (plates 3-L, 3MA, and 3HA), biotinylated oligonucleotides (200nM), DNA scaffold (20 nM final concentration), and fluorophore-labeled oligonucleotides (200 nM final concentration) were mixed in 1x folding buffer (5 mM Tris pH 8.0, 1 mM EDTA, 5 mM NaCl, 20 mM MgCl_2_). Origami folding reaction was performed in a PCR thermocycler (Bio-Rad MJ Research PTC-240 Tetrad), with initial denaturation at 65 °C for 15 min followed by cooling from 60°C to 40°C with a decrease of 1° C per hr. To purify excess oligonucleotides from fully folded DNA origami, the DNA folding reaction was mixed with an equal volume of PEG precipitation buffer (15% (w/v) PEG-8000, 5 mM Tris-Base pH 8.0, 1 mM EDTA, 500 mM NaCl, 20 mM MgCl_2_) and centrifuged at 16,000x rcf for 25 min at room temperature. The supernatant was removed, and the pellet was resuspended in 1x folding buffer. PEG purification was repeated a second time and the final pellet was resuspended at the desired concentration in 1x folding buffer and stored at 4°C.

### Preparation of benzylguanine-conjugated DNA oligonucleotides

5’-amine modified (5AmMC6) DNA oligonucleotides were ordered from IDT and diluted in 0.15 M HEPES pH 8.5 to a final concentration of 2 mM. N-hydroxysuccinimide ester (BG-GLA-NHS) functionalized benzylguanine was purchased from NEB (Cat #S9151S) and freshly reconstituted in DMSO to a final concentration of 83 mM. To functionalize the oligonucleotides with benzylguanine, the two solutions were mixed so that the molar ratio of oligonucleotide-amine:benzylguanine-NHS is 1:50, and the final concentration of HEPES is between 50 mM and 100 mM. The reaction was left on a rotator overnight at room temperature. To remove excess benzylguanine-NHS ester, the reaction product was purified the next day with illustra NAP-5 Columns (Cytiva, Cat #17085301), using H_2_O for elution. The molar concentration of the benzylguanine conjugated oligonucleotides was determined by measuring the absorbance of the purified reaction at 260 nm with a Nanodrop. This reaction was further condensed with the Savant SpeedVac DNA 130 Integrated Vacuum Concentrator System, resuspended in water to a final concentration of 100 µM, aliquoted, and stored at −20°C until use.

### Functionalization of glass surface with DNA origami

96-well glass bottom MatriPlates were purchased from Brooks (Catalog # MGB096-1-2-LG-L). Before use, plates were incubated in 5% (v/v) Hellmanex III solution (Z805939-1EA; Sigma) overnight, washed extensively with Milli-Q water, dried under the flow of nitrogen gas, and covered with sealing tape (ThermoFisher, Cat # 15036). Wells used for experiment were unsealed, incubated with 200 µL of Biotin-BSA (ThermoFisher, Cat # 29130) at 0.5 mg/mL in PBS pH 7.4 at RT for 2 hr-overnight. Wells were washed 6x with PBS pH 7.4 to remove excess BSA and incubated for 30 min at room temperature with 100 μL neutravidin at 250 μg/mL in PBS pH 7.4 for origami quantification and 50 μg/mL for cellular experiments. Wells were again washed 6x with PBS pH 7.4 supplemented with 20 mM MgCl_2_ and incubated for 1-2 hr with the desired amount of DNA origami diluted in PBS pH 7.4 with 20 mM MgCl_2_ and 0.1% BSA.

### DNA origami quantification

5 wells of a 96-well glass bottom MatriPlate per origami reaction were prepared as described in ‘Functionalization of glass surface with DNA origami’. The purified DNA origami reaction was serially diluted into PBS pH 7.4 with 20 mM MgCl_2_ and 0.1% BSA and 5 different concentrations were plated and incubated for 1.5 hr before washing 5x with PBS pH 7.4 with 20 mM MgCl_2_ and 0.1% BSA. Fluorescent TIRF images were acquired in the channel with which the origami was labeled. 100 sites per well were imaged using the High Content Screening (HCS) Site Generator plugin in uManager (Stuurman, Edelstein, Amodaj, Hoover, & Vale, 2010). The number of individual DNA origami per um^2^ in each well were quantified using the Spot Counter plugin in Fiji. This was repeated for all concentrations of origami plated. The final concentration of the origami reaction was measured as number of origami/µm^2^ and was calculated from a linear fit including all concentrations in which individual origami could be identified by the plugin.

### TIRF imaging

96-well glass bottom MatriPlates were functionalized with DNA origami as described and then washed into engulfment imaging media (20 mM Hepes pH 7.4, 135 mM NaCl, 4 mM KCl, 1 mM CaCl_2,_ 10 mM glucose) containing 20 mM MgCl_2_. ∼100,000 dual infected mNeonGreen-DNA-CAR*γ* and BFP-Syk THP1 cells per well were pelleted via centrifugation, washed into engulfment imaging media, re-pelleted, and resuspended into 50 µL of engulfment imaging media. 1uL of 100 μM benzylguanine-labeled receptor DNA stock was added per ∼50,000 cells pelleted, and the cell-DNA mixture was incubated at room temperature for 15 min. Cells were subsequently washed twice via centrifugation with 10 mL of imaging buffer to remove excess benzylguanine labeled DNA and resuspended in 200 μL per 100,000 cells of imaging buffer containing 20 mM MgCl_2_. Cells were then immediately added to each well and imaged. Data was only collected from a central ROI in the TIRF field. The origami fluorescent intensities along the x and y axis were plotted to ensure there was no drop off in signal and thus no uniformity of illumination.

### Quantification of receptor and Syk recruitment to individual origami

Cells that expressed both the mNeonGreen tagged DNA-CAR*γ* receptor and the BFP-tagged Syk and had interactions with the 72-ligand origami were chosen for analysis in Fiji. An ROI was drawn around the perimeter of the cell-glass surface interaction, which was determined by the presence of receptor fluorescence. The ‘Spot Intensity in All Channel’ plugin in Fiji was used to identify individual origami pegboards, measure fluorescence intensity of the DNA-CAR*γ* receptor and Syk at each origami pegboard, and subtract local background fluorescence. The intensity at each origami pegboard was normalized to the average intensity measured at 72-ligand origami pegboards in each well.

### Supported lipid bilayer coated silica bead preparation

Chloroform-suspended lipids were mixed in the following molar ratios: 96.8% POPC (Avanti, Catalog # 850457), 2.5% biotinyl cap PE (Avanti, Catalog # 870273), 0.5% PEG5000-PE (Avanti, Catalog # 880230, and 0.2% atto390-DOPE (ATTO-TEC GmbH, Catalog # AD 390–161) for labeled lipid bilayers, or 97% POPC, 2.5% biotinyl cap PE, and 0.5% PEG5000-PE for unlabeled lipid bilayers. The lipid mixes were dried under argon gas and desiccated overnight to remove chloroform. The dried lipids were resuspended in 1 mL PBS, pH 7.2 (Gibco, Catalog # 20012050) and stored under argon gas. Lipids were formed into small unilamellar vesicles via ≥30 rounds of freeze-thaws and cleared via ultracentrifugation (TLA120.1 rotor, 35,000 rpm / 53,227 x g, 35 min, 4°C). Lipids were stored at 4°C under argon gas in an eppendorf tube for up to two weeks. To form bilayers on beads, 8.6 × 10^8^ silica beads with a 4.89 µm diameter (10 µl of 10% solids, Bangs Labs, Catalog # SS05N) were washed 2x with water followed by 2x with PBS by spinning at 300rcf and decanting. Beads were then mixed with 1mM SUVs in PBS, vortexed for 10 s at medium speed, covered in foil, and incubated in an end-over-end rotator at room temperature for 0.5-2 hr to allow bilayers to form over the beads. The beads were then washed 3x in PBS to remove excess SUVs, and resuspended in 100uL of 0.2% casein (Sigma, catalog # C5890) in PBS for 15 min at room temperature to block nonspecific binding. Neutravidin (Thermo, Catalog # 31000) was added to the beads at a final concentration of 1 ug/ml for 20-30 minutes, and the beads were subsequently washed 3x in PBS with 0.2% casein and 20mM MgCl_2_ to remove unbound neutravidin. The indicated amounts of biotinylated ssDNA or saturating amounts of DNA origami pegboards were added to the beads and incubated for 1 hr at room temperature with end-over-end mixing to allow for coupling. Beads were washed 2 times and resuspended in 100uL PBS with 0.2% casein and 20 mM MgCl_2_ to remove uncoupled origami pegboards or ssDNA. When functionalizing SUV-coated beads with anti-biotin AlexaFluor647-IgG (Jackson ImmunoResearch Laboratories Catalog # 200-602-211, Lot # 137445), the IgG was added to the beads at 1uM immediately following the casein blocking step, and beads were incubated for 1 hr at room temperature with end-over-end mixing.

### Quantification of ssDNA, IgG, or origami on beads

To estimate the amount of ssDNA bound to each bead, we compared the fluorescence of Atto647-labeled DNA on the bead surface to calibrated fluorescent beads (Quantum AlexaFluor 647, Bangs Lab) using confocal microscopy (Figure S1). To determine saturating conditions of IgG and origami pegboards, we titrated the amount of IgG or origami in the coupling reaction and used confocal microscopy to determine the concentration at which maximum coupling was achieved. A comparable amount of origami pegboard coupling was also confirmed with confocal microscopy for beads used in the same experiment.

### Quantification of engulfment

30,000 RAW264.7 macrophages were plated in one well of a 96-well glass bottom MatriPlate (Brooks, Catalog # MGB096-1-2-LG-L) between 12 and 24 hr prior to the experiment. Immediately before adding beads, 100 μL of a 1 μM solution of benzylguanine-conjugated receptor DNA in engulfment imaging media was added, incubated for 10 min at room temperature, and washed out 4 times with engulfment imaging media containing 20 mM MgCl_2_, making sure to leave ∼100 μL of media covering the cells between washes, and finally leaving the cells in ∼300 μL of media. ∼8 × 10^5^ beads were added to the well and engulfment was allowed to proceed for 45 min in the cell incubator. Cells were fixed with 4% PFA for 10 min and washed into PBS. For figures 4c and 6d, 10 nM AlexaFluor647 anti-biotin IgG (Jackson Immuno Labs, Catalog # 200-602-211) diluted into PBS containing 3% BSA was added to each well for 10 minutes to label non-internalized beads. Wells were subsequently washed 3 times with PBS. Images were acquired using the High Content Screening (HCS) Site Generator plugin in µManager and at least 100 cells were scored for each condition. When quantifying bead engulfment, cells were selected for analysis based on a threshold of GFP fluorescence, which was held constant throughout analysis for each individual experiment. For figures 3, 4, 6, and S5 the analyzer was blinded during engulfment scoring using the position randomizer plug-in in µManager. For the THP1 cells, ∼100,000 cells per condition were spun down, washed into engulfment imaging media, and coupled to benzylguanine-labeled receptor DNA as described under TIRF imaging. Cells were resuspended into 300 μL engulfment imaging media containing 20 mM MgCl_2_ in an Eppendorf tube, ∼8 × 10^5^ beads were added to the tube, and the tube was inverted 8x before plating the solution into a round-bottomed 96 well plate (Corning, Catalog # 38018). Engulfment was allowed to proceed for 45 min in the cell incubator before the plate was briefly spun and the cells were fixed in 4% PFA for 10 min. Cells were subsequently washed 3x with PBS by briefly centrifuging the plate and removing the media, and finally moved into a 96-well glass bottom MatriPlate for imaging.

### Quantification of engulfment kinetics

RAW264.7 macrophages were plated and prepared in wells of a 96-well glass bottom MatriPlate as described in ‘Quantification of engulfment’. Using Multi-Dimensional Acquisition in µManager, 4 positions in the well were marked for imaging at 20 sec intervals through at least 7 z-planes. ∼4 × 10^5^ Atto647N-labeled 4S origami functionalized beads and ∼4 × 10^5^ Atto550N-labeled 4T origami functionalized beads were mixed in an Eppendorf tube, added to the well, and immediately imaged. Bead contacts were identified by counting the number of beads that came into contact with the cells throughout the imaging time. Initiation events were identified by active membrane extension events around the bead. Engulfment completion was identified by complete internalization of the bead by the macrophage. The initiation time was quantified as the amount of time between bead contact (the first frame in which the bead contacted the macrophage) and engulfment initiation (the first frame in which membrane extension around the bead was visualized) and was only measured for beads that were completely internalized by the end of the imaging time. The engulfment time was quantified as the amount of time between engulfment initiation and engulfment completion (the first frame in which the bead has been fully internalized by the cell).

### Quantification of synapse intensity of DNA-CAR*γ* receptor, tSH2 Syk, PIP_3_ reporter, and actin filaments

Phagocytic cups were selected for analysis based on clear initiation of membrane extension around the bead visualized by GFP fluorescence from the DNA-CAR*γ* receptor. The phagocytic cup and the cell cortex (areas indicated in schematic in figure 6b) were traced with a line (6 pixels wide for DNA-CAR*γ* receptor and the tSH2 Syk reporter, and 8 pixels wide for the Akt-PH reporter and phalloidin) at the Z-slice with the clearest cross section of the cup.

### Microscopy and analysis

Images were acquired on a spinning disc confocal microscope (Nikon Ti-Eclipse inverted microscope with a Yokogawa CSU-X spinning disk unit and an Andor iXon EM-CCD camera) equipped with a 40 × 0.95 NA air and a 100 × 1.49 NA oil immersion objective. The microscope was controlled using µManager. For TIRF imaging, images were acquired on the same microscope with a motorized TIRF arm using a Hamamatsu Flash 4.0 camera and the 100x 1.49 NA oil immersion objective.

### Statistics

Statistical analysis was performed in Prism 8 (GraphPad, Inc). The statistical test used is indicated in each relevant figure legend.

## Acknowledgements

We thank N. Stuurman for help with microscopy and developing the ‘image randomizer’ plug-in for blinding our analysis as well as the ‘Spot Intensity in All Channel’ plugin for quantification of our TIRF experiments. We also thank K. McKinley, T. Skokan, C. Gladkova, J. Sheu-Gruttadauria for discussions and critical feedback on this manuscript. M.A.M. was supported by the National Institute of General Medical Sciences of the National Institutes of Health under award number F32GM120990. S.M.D. was supported by the Army Research Office (W911NF-14-1-0507) and Office of Naval Research (N00014-17-1-2627). Funding was provided by the Howard Hughes Medical Institute to R.D.V.

## Author Contributions

N.K., R.D.V., and M.A.M. designed research; N.K. performed research; N.K., R.D., S.M.D. and M.A.M. contributed new resources; N.K. analyzed data; and N.K., R.D.V., S.M.D., and M.A.M wrote the paper.

