## Supplementary figures and images for "Tight nanoscale clustering of Fcγ-receptors using DNA origami promotes phagocytosis"

### Supplemental Figures

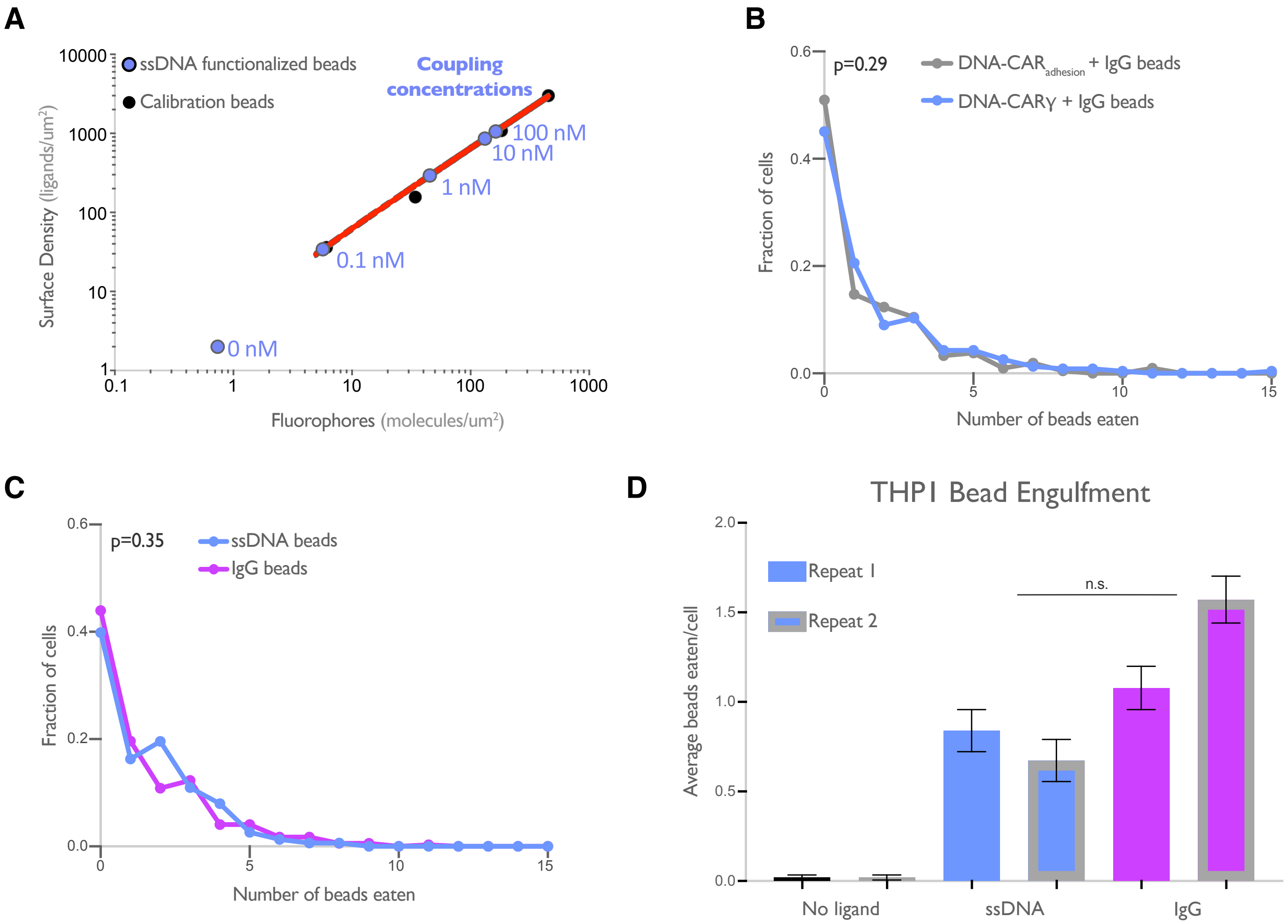

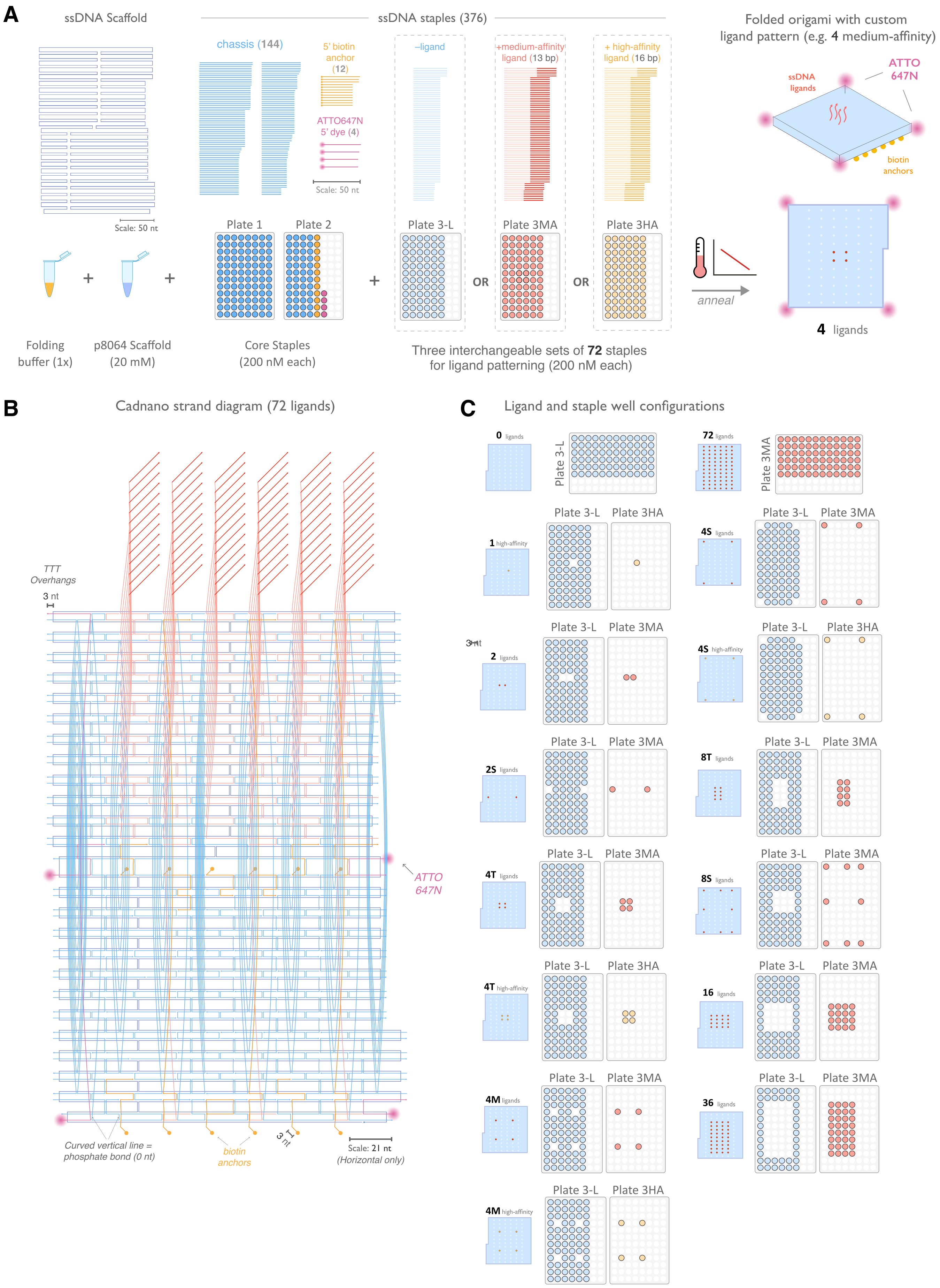

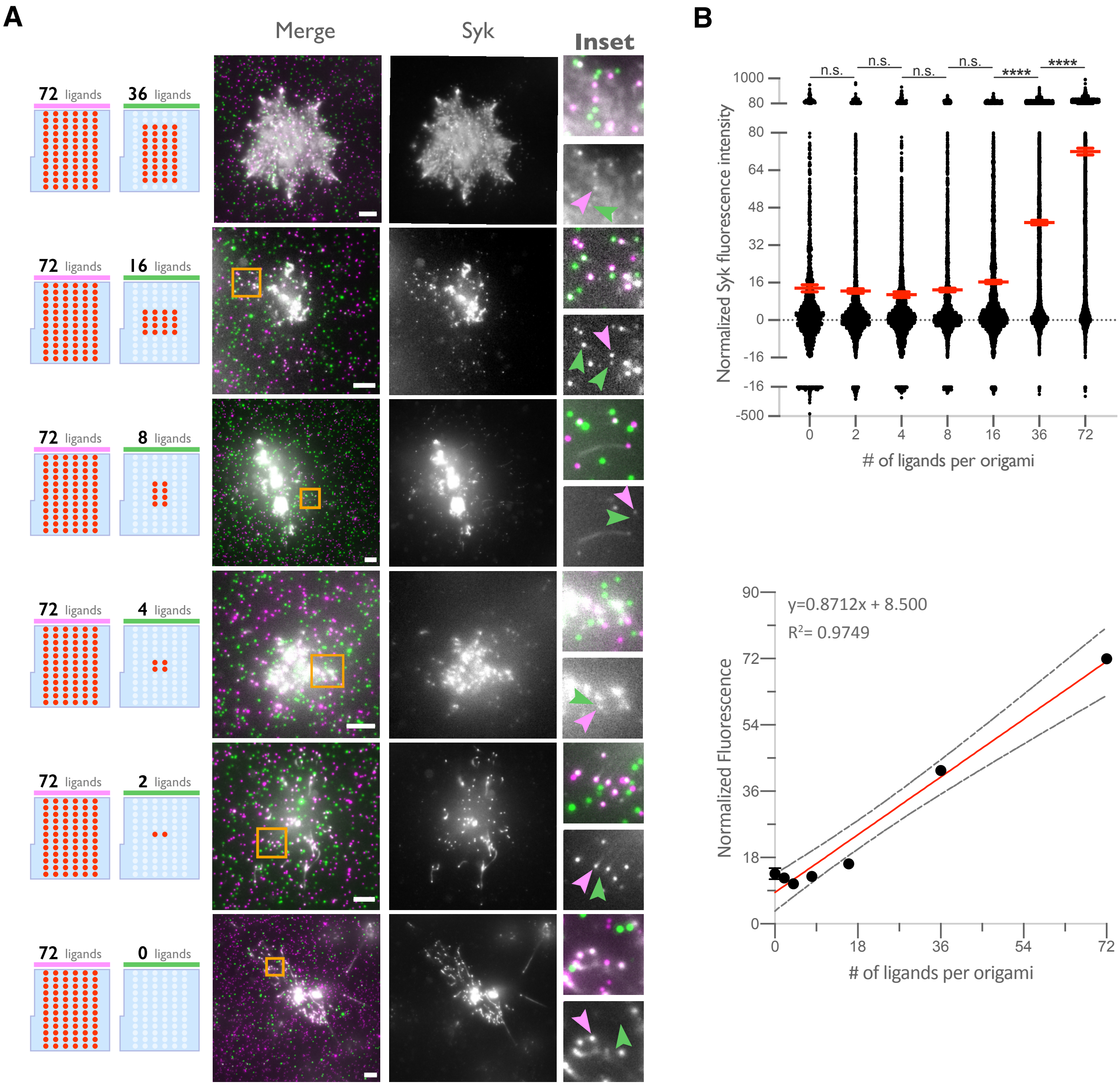

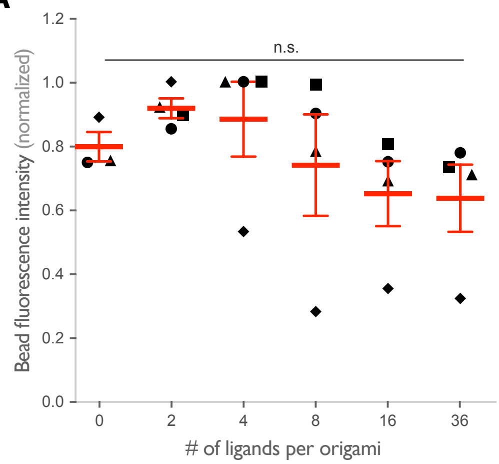

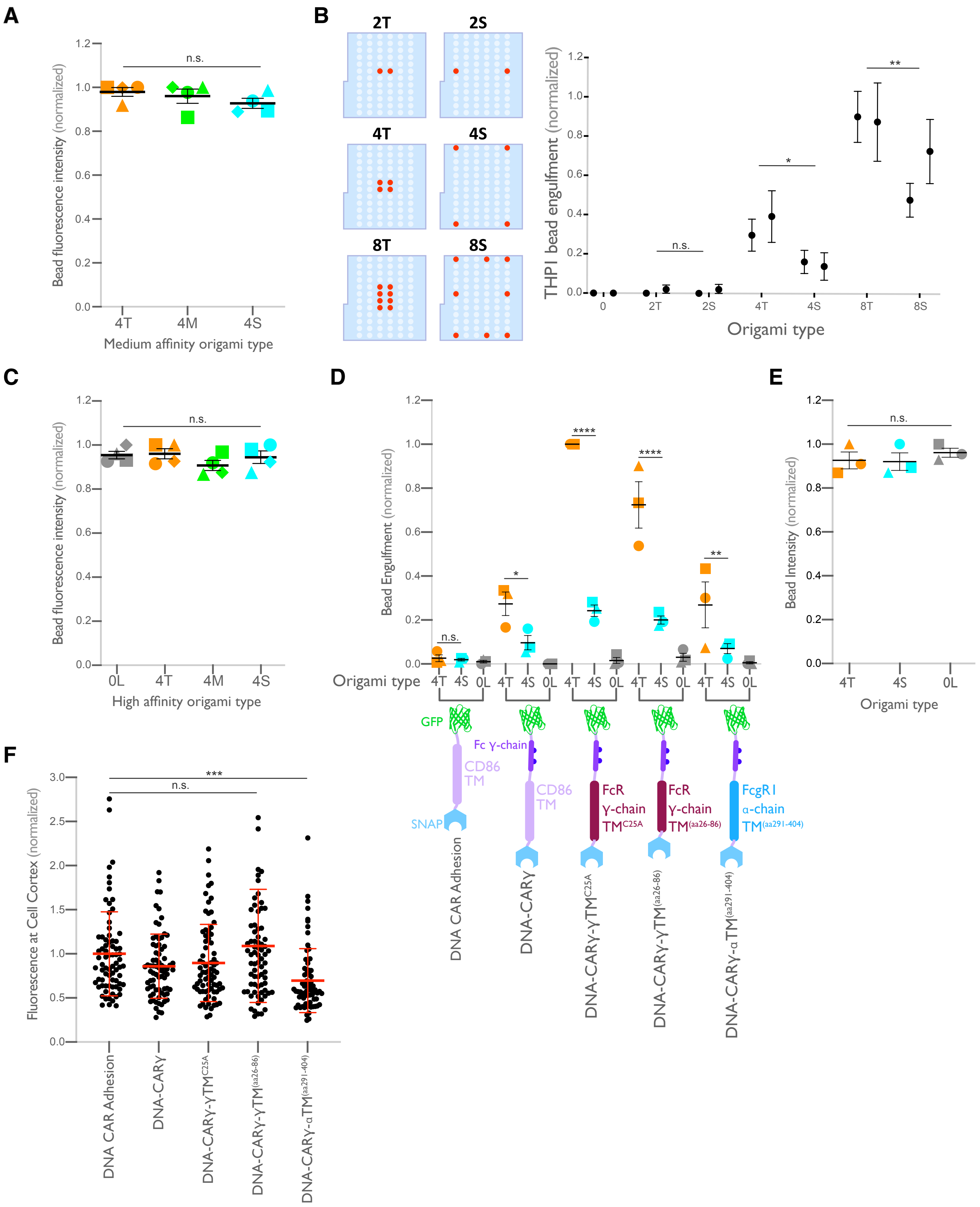

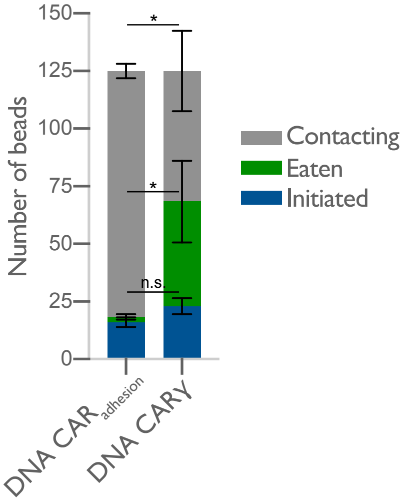

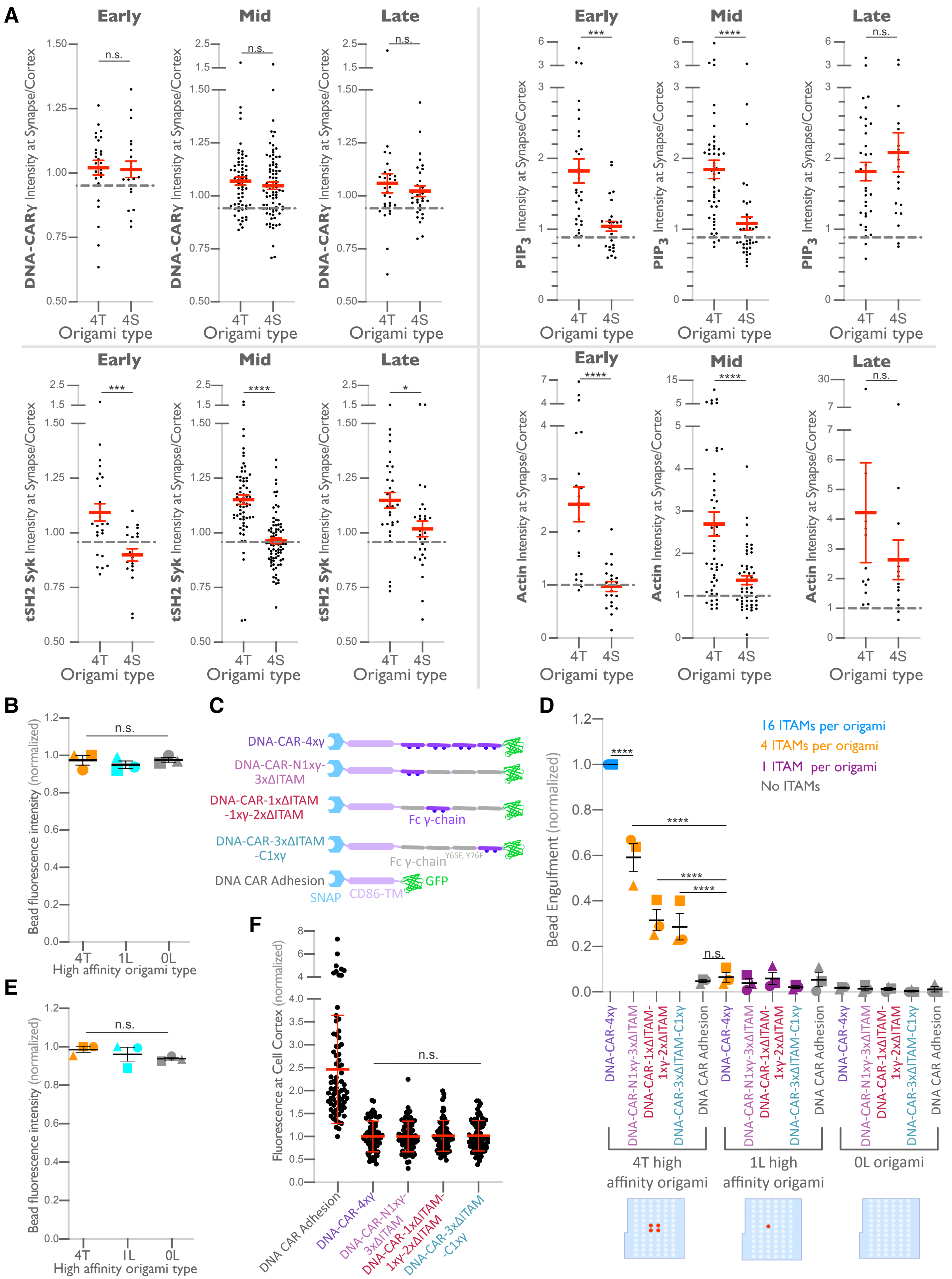
